# Mechanism of D-type cyclins recognition by the AMBRA1 E3 ligase receptor

**DOI:** 10.1101/2024.12.17.628619

**Authors:** Yang Wang, Ming Liu, Shan Wang, Xinyi Mai, Xi Wang, Fei Teng, Tianrui Lyu, Ming-Yuan Su, Goran Stjepanovic

## Abstract

AMBRA1 is a tumour suppressor protein that functions as a substrate receptor in the ubiquitin conjugation system and regulates the stability of D-type cyclins and cell proliferation. Here, we present the cryo-EM structure of cyclin D1 bound AMBRA1-DDB1 complex at 3.5 Å resolution. The structure reveals a substrate interaction surface on the AMBRA1 WD40 domain that specifically binds to the C-terminal region of D-type cyclins. This interaction is dependent on the phosphorylation of Thr286 residue in the C-terminal phosphodegron site of D-type cyclins. The phosphodegron motif folds into a turn-like conformation followed by a 3_10_ helix that promotes its assembly with AMBRA1. Additionally, we show that AMBRA1 mutants, which are defective in cyclin D1 binding, lead to cyclin D1 accumulation and DNA damage. Understanding the AMBRA1-D-type cyclins structure enhances the knowledge of the molecular mechanisms that govern the cell cycle control and may lead to new therapeutic approaches for cancers linked to abnormal cyclin D activity.

## Introduction

The ubiquitin-proteasome system (UPS) is a highly regulated and intricate cellular machinery responsible for maintaining protein homeostasis and controlling key cellular processes. It involves a series of enzymatic reactions that result in the attachment of ubiquitin molecules to target proteins, marking them for degradation by the proteasome. The UPS plays a crucial role in removing aberrant or damaged proteins that could be harmful to the cell. Additionally, it regulates the turnover of numerous proteins involved in various cellular functions, such as cell cycle progression, signal transduction, and protein quality control. One specific aspect of cell cycle regulation where the UPS is highly involved is the control of cyclin levels and cyclin-dependent kinases (CDKs) activity (*1*). Cyclins are proteins that associate with and activate CDKs at specific points in the cell cycle. In particular, the D-type cyclins, which are encoded by three closely related genes (cyclins D1, D2, and D3), play a critical role in responding to extracellular mitogenic signals and govern progression through G1 phase via their association with CDK4 and CDK6. Proteasome-mediated degradation serves as a critical mechanism for regulating the steady-state levels of cyclin D1 and ensuring the proper control of cell cycle progression. When cyclin D1 is deregulated, either through overexpression, accumulation, or inappropriate localization, it becomes an oncogene and contributes to genomic instability and tumour development (*2*). Aberrant overexpression of cyclin D1 is frequently observed in human cancers (*3*). This deregulation serves as a biomarker of the cancer phenotype and progression of the disease (*4, 5*). Cyclin D1 and its associated CDKs are promising therapeutic targets in human cancers. However, the precise mechanisms that regulate cyclin D degradation and cellular levels are still not fully understood.

Central to the UPS is the action of ubiquitin E3 ligases, which are enzymes responsible for recognizing specific protein substrates and facilitating the transfer of ubiquitin molecules onto them. Within eukaryotes, the Cullin-RING ligases (CRLs) constitute the largest family of E3 ligases. These ligases are composed of a scaffold protein Cullin and a catalytic RING subunit, RBX1 or RBX2. The Damage-specific DNA binding protein 1 (DDB1) acts as an integral component of the Cullin4A/B-RING E3 ubiquitin ligase (CRL4) complex. Its serves as an adaptor protein between Cullin4A/B (CUL4A/B) and CUL4A-associated factors (DCAFs), facilitating the targeting of substrates for ubiquitination. DDB1 possesses three β-propeller domains (BPA, BPB, and BPC) and has the ability to interact with various substrate receptors. These receptors, in turn, recruit specific substrates for further processing.

AMBRA1 (Autophagy and Beclin 1 Regulator 1) functions as E3 ligase receptor involved in the regulation of autophagy and cell cycle control. AMBRA1 acts as a substrate recognition subunit of the CRL4^DDB1^ E3 ligase complex and facilitates the ubiquitination and proteasomal degradation of its target proteins. AMBRA1 functions as the main regulator of the degradation of D-type cyclins (Fig. 1A) (*6–8*). AMBRA1 recognizes and ubiquitinates the C-terminal phosphodegron of D-type cyclins that contains a highly conserved threonine at position 286 (*1, 7, 9*). Phosphorylation of Thr286 by glycogen synthase kinase 3β facilitates the binding of cyclin D1 with the nuclear exportin CRM1, and triggers cyclin D1 nuclear export and proteolytic turnover (*10, 11*). Mutation that blocked the phosphorylation of Thr286 (T286A) was shown to inhibit cyclin D1 degradation via UPS in mouse fibroblasts (*9*). Similarly, T286A mutation impaired AMBRA1-cyclin D1 interaction and stabilized cyclin D1 in human cell lines (*7*). Thr286 is mutated in a variety of human cancers, and overexpression of mutant alleles in animal models drives spontaneous tumours (*1, 12*). AMBRA1 is a tumour suppressor that is mutated in a wide range of human tumours (*13, 14*). Loss of AMBRA1 and the resulting stabilization of cyclin D1 leads to increased progression of the cell cycle, genomic instability and decreased sensitivity to CDK4/6 inhibitors (*7, 8, 15, 16*). Therefore, AMBRA1 has emerged as a novel target for anticancer treatment and biomarker for cancer therapy (*17*).

**Fig. 1.**
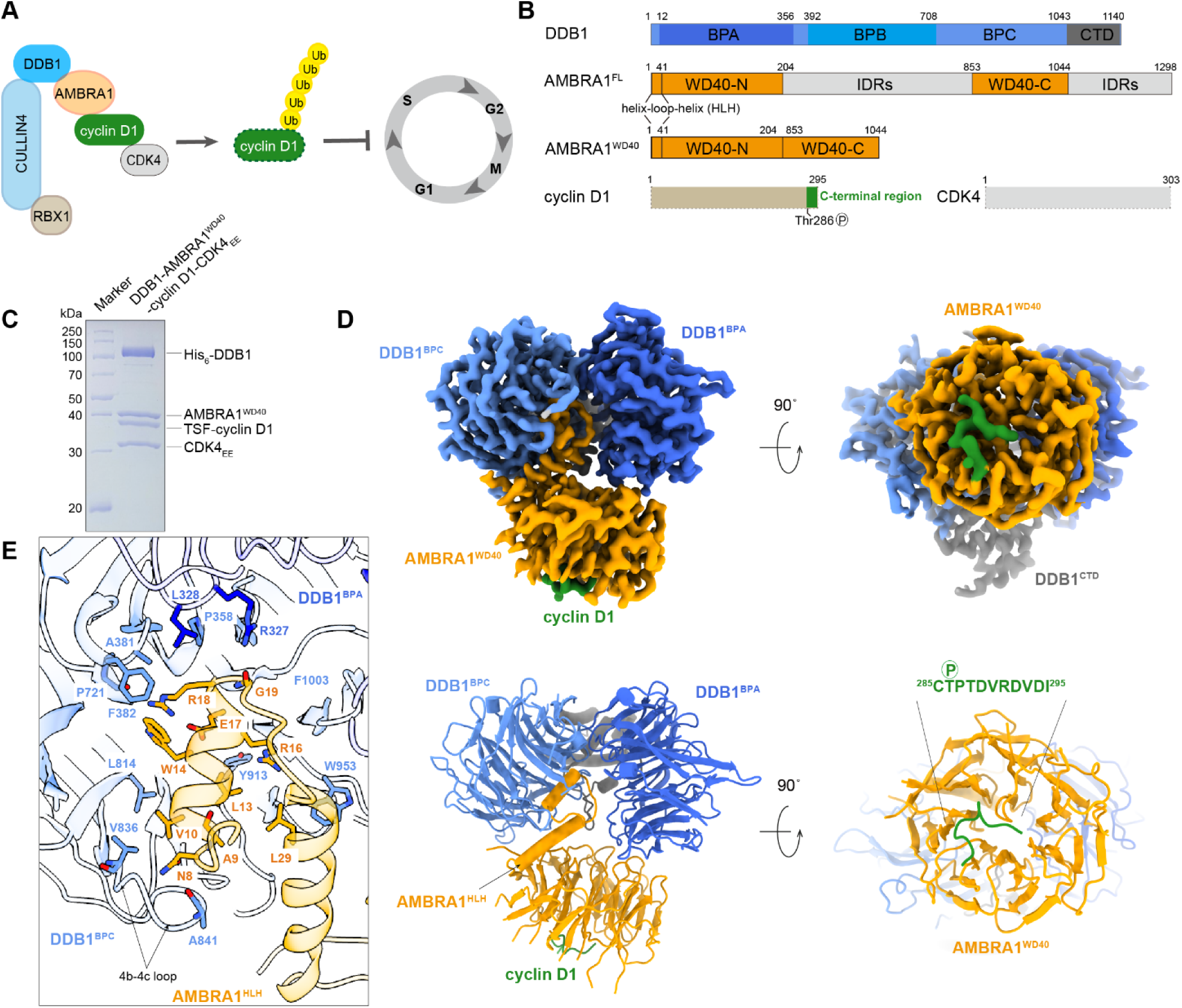
The cryo-EM structure of DDB1-AMBRA1^WD40^-cyclin D1. **(A)** Cartoon schematic of Cullin4-DDB1-RBX1-AMRBA1 E3 ligase mediating ubiquitination of cyclin D1 in regulating cell cycle progression. **(B)** Annotated DDB1, AMBRA1, cyclin D1 and CDK4 domain schematics. The AMBRA1 ^WD40^ construct includes the N-terminal helix-loop-helix, WD40-N and WD40-C. BPA-BPC, β-propeller A-C; CTD, C-terminal helical domain; FL, full-length; IDRs, intrinsically disordered regions. **(C)** Coomassie blue-stained SDS-PAGE analysis of purified DDB1-AMBRA1 ^WD40^-cyclin D1- CDK4_EE_ complex used for cryo-EM structure determination. TSF, twin-strep-flag. **(D)** The cryo-EM density map and the refined coordinates of the DDB1- AMBRA1 ^WD40^-cyclin D1 complex. AMBRA1^WD40^ is colored in orange; BPA domain in dark blue; BPC-CTD domain in light blue and gray; C-terminal region of cyclin D1 in green. **(E)** Close-up view of the AMBRA1^HLH^-DDB1 interface. The key residues contributing to the interface are labelled.

Recent studies showed that AMBRA1 is a highly dynamic and largely intrinsically disordered protein. AMBRA1 contains a “split” WD40 domain (AMBRA1^WD40^) consisting of two halves separated by a long, intrinsically disordered region (Fig. 1B). These two parts need to reunite to form a seven-bladed β-propeller that binds to DDB1 through the N-terminal helix-loop-helix motif (*15, 18*). This interaction was shown to be similar to other E3 substrate receptors bound to DDB1, including DCAF1, Cockayne syndrome group A protein (CSA) and Simian virus 5 V (*15*). We have previously shown that AMBRA1 WD40 domain is sufficient for cyclin D1 recognition and ubiquitination, highlighting its functional role in substrate recruitment (*15*). However, the molecular mechanism of substrate binding by AMBRA1 is unknown. In addition to cyclin D1, numerous other AMBRA1 interacting proteins have been identified (*19*), allowing AMBRA1 to function as a hub in various physiological processes, including autophagy, development, cell death and proliferation. Structures of AMBRA1 in complex with substrates are necessary to provide insight into the recognition of diverse target proteins and set a foundation for development of new therapeutic strategies.

Here, we report the cryo-EM structure of the human DDB1-AMBRA1^WD40^ E3 ligase receptor complex bound to the cyclin D1-CDK4 complex, as a substrate. The structure reveals a protein-protein interacting surface on AMBRA1^WD40^ that binds to the extreme C-terminus of cyclin D1 and this interaction is dependent on phosphorylation at Thr286. Experimental structures allowed for comparison of AlphaFold3 models and *in silico* analysis of AMBRA1 binding to other D-type cyclins. The *in vitro* biochemical studies show that the AMBRA1 WD40 domain can bind and mediate ubiquitination of the cyclin D1 C-terminus in a phosphorylation-dependent manner. Consistent with these data, we further show that mutations that disrupt the AMBRA1-cyclin D1 interaction cause endogenous cellular DNA damage. These studies provide important mechanistic insights into the AMBRA1-cyclin D1 recognition and suggest a common mechanism for AMBRA1-mediated ubiquitination of D-type cyclins in a phosphorylation-dependent manner.

## Results

### Structure of DDB1-AMBRA1^WD40^ bound to cyclin D1-CDK4

To understand AMBRA1 substrate recognition, we determined the structure of the cyclin D1-CDK4 in complex with AMBRA1^WD40^ domain and DDB1 (DDB1-AMBRA1^WD40^) by single-particle Cryo-EM at a resolution of 3.5Å. The protein complex was purified from Expi293F cells by coexpressing human AMBRA1 WD40 domain with full-length cyclin D1, CDK4_EE_ and DDB1. In the CDK4 construct, amino acids G43-G47 were replaced with EE (CDK4_EE_) in order to increase the protein stability (*20*). AMBRA1^WD40^ construct was generated as described previously, and comprised residues M1-N204 directly fused to residues S853-G1044 (*15*). The purified DDB1- AMBRA1^WD40^-cyclin D1-CDK4_EE_ complex appeared as a stable heterotetramer judged by SDS‒ PAGE and size exclusion chromatography (Fig. 1C, Fig. S1A). Cryo-EM images were collected and processed as detailed in the “Materials and Methods” section (Fig. S2). 2D classification and 3D refinement procedures converged to the part of the complex containing DDB1 and AMBRA1^WD40^. The resulting electron density map allowed unambiguous tracing of the AMBRA1^WD40^ domain, BPA and BPC domains of DDB1, and the extreme C-terminus of cyclin D1, allowing us to build a reliable atomic model (Fig. 1D). Post-translational modification analysis including western blotting and mass spectrometry of the reconstituted protein complex which was used for structural determination revealed the presence of Thr286 phosphorylation in cyclin D1 (Fig. S1H, I).

The overall structure of the DDB1-AMBRA1^WD40^ receptor subcomplex agrees well with our previous reported structure without the substrate (*15*), with the map and model quality substantially improved. Interaction between AMBRA1 and DDB1 is mediated by helix-loop-helix motif located at the N-terminus of AMBRA1 (residues E6-K41). This motif inserts in the binding pocket formed between BPA and BPC double propeller of DDB1 (Fig. 1D, E). The side chain of N8 and the backbone NH of A9, located at the N-terminal end of the first helix of AMBRA1, form hydrogen bonds with the backbone CO groups in the loop connecting β-strands b and c in blade IV of the DDB1 BPC domain. The side chains of residues A9, V10, L13, and W14 in AMBRA1 contribute to the interactions by packing against the surface of the DDB1 BPC domain. At the C-terminal end of the first helix, the side chains of residues R16 and R18 form multiple hydrogen bonds with DDB1. R16 forms hydrogen bonds with Y913 of DDB1 while the guanidinium group of R18 interacts with the side chain of E17 in AMBRA1 and the backbone carbonyl group of P721 in DDB1. The sidechain methylene groups of R18 and the side chain of W14 are accommodated by a hydrophobic patch on the BPC domain formed by L328, P358, A381 and F382. Finally, R327 in DDB1 forms a hydrogen bond with the backbone carbonyl group of G19 (Fig. 1E). This interaction interface is further strengthened by the second helix of AMBRA1, which packs against helix 1 and also interacts with the surface of the DDB1 BPC domain (Fig. 1E). The Cullin binding domain (BPB) of DDB1 exhibits high conformational heterogeneity and was subtracted during data processing to improve resolution of the final reconstruction (Fig. S2).

Structured regions of AMBRA1 consist of the N-terminal helix-loop-helix motif followed by the WD40 domain. The AMBRA1^WD40^ domain exhibits a β-propeller architecture, comprised of seven blades and each blade contains four anti-parallel β-strands (a-d). The AMBRA1^WD40^ β-propeller structure has a wider bottom and a narrower top when observed from the side view, and packs against the BPA and BPC domains of DDB1 with a bottom surface (Fig. 1D, 2A). One distinctive structural feature of AMBRA1 is presence of the 650-residue intrinsically disordered region in blade IV, between the b and c β-strands. This “split” WD40 domain organization of AMBRA1 is stabilized by DDB1 and proposed to facilitate the interaction between diverse substrates and CRL4 E3 ligase, thereby aiding in the transfer of ubiquitin (*15*). The density for most of the cyclin D1- CDK4_EE_ subcomplex was not visible, probably due to its flexibility, and only the last eleven residues of cyclin D1 assume well-defined conformation in the electron density map. From overall structure, the extreme C-terminal region of cyclin D1 (residues C285−I295) binds in a turn-like conformation to the top surface of AMBRA1^WD40^, and covers the central pore of the β-propeller (Fig. 1D, 2A). The binding of the substrate to the exposed top surface of AMBRA1^WD40^ orients the substrate for subsequent ubiquitination by the E3 ligase complex.

**Fig. 2.**
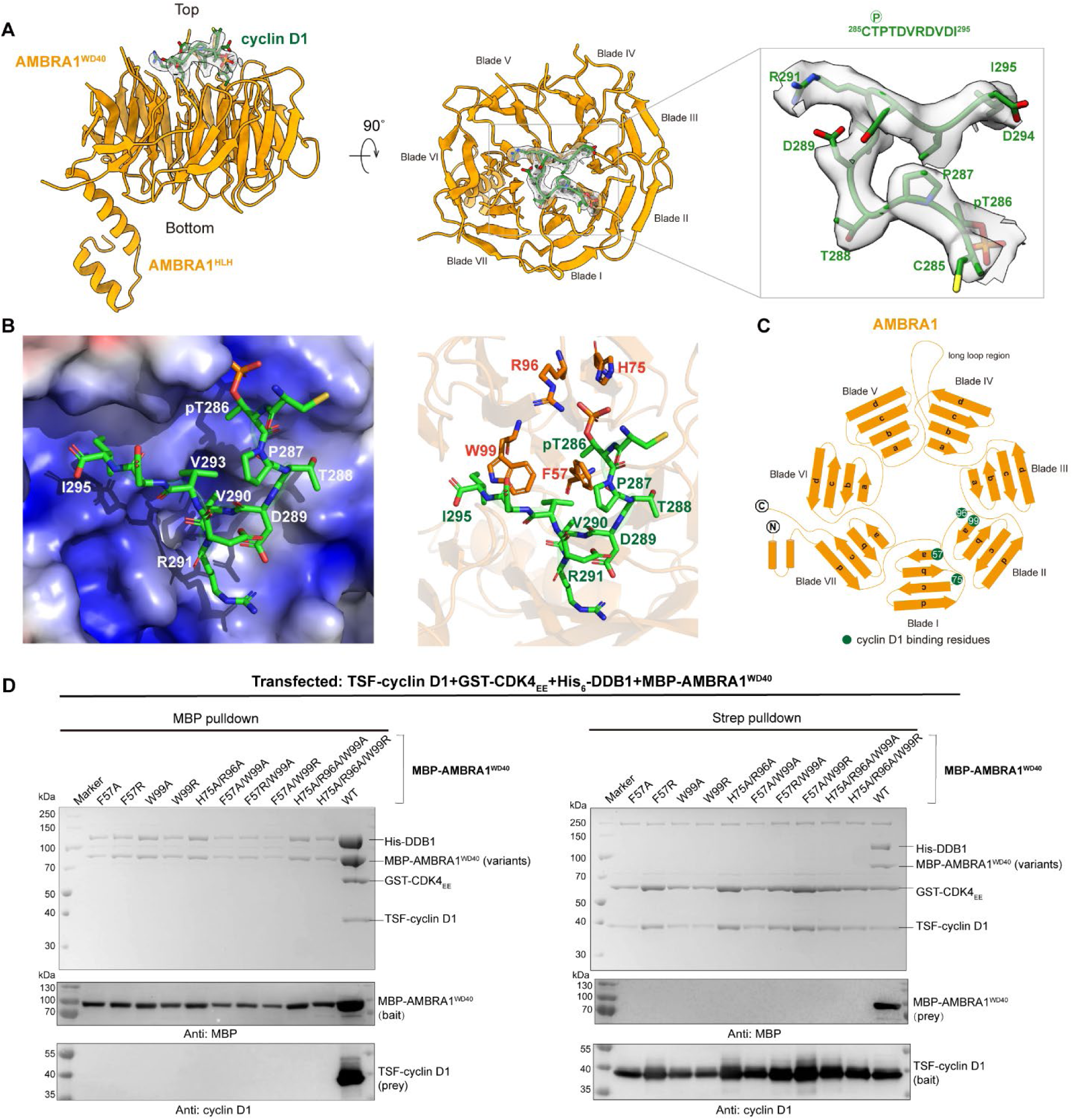
The interface of the AMBRA1 ^WD40^-cyclin D1 complex. **(A)** The model of AMBRA1^WD40^-cyclin D1 complex. Close-up view of map density and coordinate of cyclin D1 C-terminal region. **(B)** The electrostatics potential of the AMBRA1^WD40^-cyclin D1 complex interface. The key residues contributing to the interaction of AMBRA1 ^WD40^ are labelled. **(C)** The cartoon of the AMBRA1, and the key binding residues are labelled in green cycle. **(D)** *In vitro* pull-down experiments of MBP-AMBRA1^WD40^ or key binding residues mutants with TSF-cyclin D1. The cells were co-transfected with TSF-cyclin D1, GST-CDK4_EE_, His_6_-DDB1 and MBP-AMBRA1^WD40^. The experiment was repeated at least three times and visualized by SDS- PAGE and western blotting.

### Molecular interactions of AMBRA1^WD40^ with cyclin D1

The specificity of phosphorylation-dependent recognition of cyclin D1 can be entirely attributed to the structural features present within the WD40 domain of AMBRA1, and includes hydrophobic, hydrogen-bonding, and electrostatic interactions. The interaction interface between cyclin D1 C- terminus and AMBRA1^WD40^ spans across blades I-II and extends to the centre of the β-propeller. pThr286 phosphate group interacts with the side chains of R96 and H75 and form hydrogen bond with the backbone NH of T97 in AMBRA1^WD40^ (Fig. 2B, C). The residues that follow the phosphorylation site adopt a turn-like conformation which is immediately followed by a single turn 3_10_-helix formed by residues D289−D292. This structure tightly packs against the top surface of AMBRA1^WD40^ through nonpolar interactions between the side chains of residues V290, V293 and I295 in cyclin D1, and a hydrophobic pocket flanked by the side chains of residues F57 and W99 in AMBRA^WD40^. The C-terminal carboxylate group of I295 is stabilized by positive charges that outline the cyclin D1-binding site in AMBRA^WD40^ (Fig. 2B). This interaction is complemented by insertion of the side chain of I295 into a hydrophobic pocket of AMBRA1. Similar packing arrangement of the hydrophobic C-terminal residue of cyclin D1 was observed in the SCF/FBXO31-cyclin D1 complex which is involved in DNA damage-induced cyclin D1 degradation, although this interaction was phosphorylation independent (*21*). There are no large structural rearrangements in AMBRA1^WD40^ upon binding of cyclin D1, based on a root mean square deviation (RMSD) of 1.322 Å (Fig. S3A). Taken together, cyclin D1 recognition by the WD40 domain of AMBRA1 is determined by several factors, including a specific binding site for pThr, a hydrophobic binding pocket that preferentially accommodates nonpolar residues positioned downstream of the phosphorylation site, and an electrostatic surface potential that favours acidic residues/functional groups.

The key cyclin D1 binding residues in AMBRA1^WD40^ were tested by mutagenesis. The hydrophobic residues were mutated alanine or arginine, and the positively charged amino acids were replaced with alanine. The interaction between cyclin D1 and AMBRA1^WD40^ was tested by pull-down analysis. We co-expressed the Twin-Strep-FLAG (TSF)-cyclin D1, GST-CDK4_EE_, His-DDB1 and MBP-AMBRA1^WD40^ variants harbouring mutations in the pThr286 binding site and the hydrophobic pocket, including: F57A, F57R, W99A, W99R, H75A/R96A, F57A/W99A, F57R/W99A, F57A/W99R, H75A/R96A/W99A and H75A/R96A/W99R. The MBP pull-down demonstrated that none of the AMBRA1^WD40^ mutants was able to bind cyclin D1. Similar results were shown in Strep pull-down, only cyclin D1 was able to bind to the wild-type AMBRA1^WD40^ (Fig. 2D). 2D class averages of purified AMBRA1 F57A, W99R and H75A/R96A mutants in complex with DDB1 showed the entirety of the AMBRA1^WD40^ domain and are similar to the wild-type protein, indicating that point mutations did not cause protein misfolding or influence interaction with DDB1 (Fig. S4). In summary, the pull-down experiments confirmed that the AMBRA1^WD40^ interacts with cyclin D1 C-terminus through the residues F57, H75, R96 and W99, and that both the pT268 binding site and the hydrophobic pocket are essential for the intermolecular interaction.

Earlier studies have shown that CRL4^AMBRA1^ complex can target cyclin D1, cyclin D2 and cyclin D3 for proteasome-mediated degradation (*22*). All three D-type cyclins exhibit a high degree of sequence and domain conservation, including the C-terminal threonine phosphodegron site (*1, 16*). Thr286 and the downstream hydrophobic amino acids that directly interact with AMBRA^WD40^ are conserved in cyclin D2 and cyclin D3 (Fig. 3A). An AlphaFold3 (*23*) predicted model of the human AMBRA1^WD40^-cyclin D1 complex agrees well with our experimental structure with a RMSD of 0.324 Å for the cyclin D1 C-terminus (Fig. S3B-D). This result prompted us to perform *in silico* analysis of AMBRA1 interaction with the phosphodegron sequences of cyclin D2 and cyclin D3. The AlphaFold3 predicted models show that the C-termini of cyclin D2 and cyclin D3 adopt the same turn-like conformation which is followed by a 3_10_-helix, and bind to the top surface of AMBRA1^WD40^ in a manner nearly identical to cyclin D1 (Fig. 3B-E). pThr interaction mode with AMBRA1^WD40^ is identical in all three models. Therefore, the broad structural features described here are consistent with the AMBRA1 role as a major regulator of cyclin D levels in cells, and underscore the key role of phosphorylation in regulating cyclin D recognition and degradation via the ubiquitin–proteasome pathway.

**Fig. 3.**
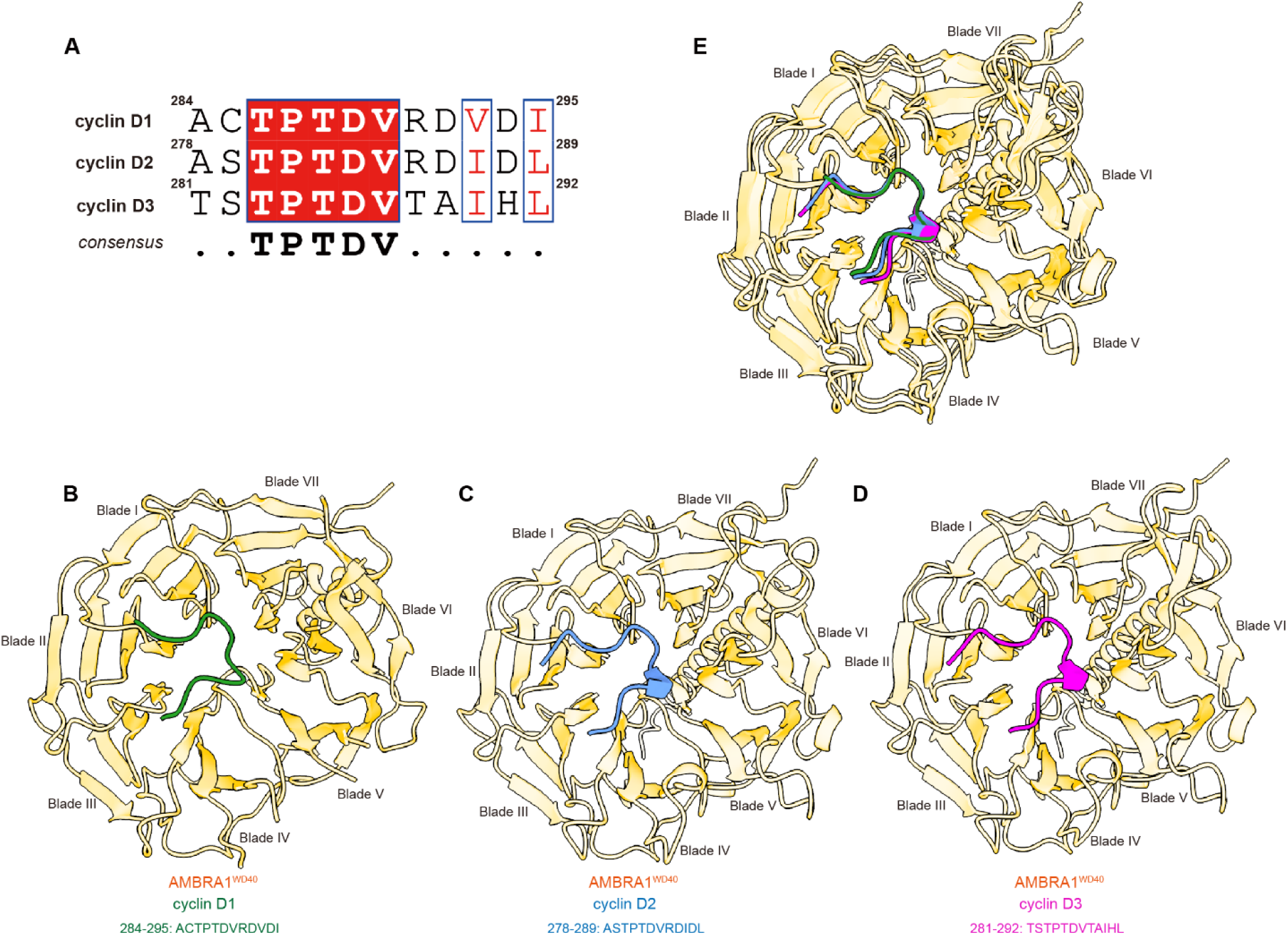
Comparison of D-type cyclins in complex with AMBRA1^WD40^. **(A)** The sequence alignment of the D-type cyclins. Cyclin D1 peptides used for alignment are residues A284-I295 resolved in cryo-EM structure, and its corresponding cyclin D2 residues A278– L289 and cyclin D3 residues T281–L292. Residues conserved among D-type cyclins are highlighted in Red. The experimental model AMBRA1^WD40^-cyclin D1 (**B**), AlphaFold3 predicted model of AMBRA1^WD40^-cyclin D2 (**C**) and AMBRA1^WD40^-cyclin D3 (**D**). **(E)** The overlap of the D-type cyclins and AMBRA1^WD40^ complex.

### AMBRA1^WD40^ recognizes C-terminal phosphodegron of D-type cyclins and mediate their ubiquitination

To address whether cyclin D1 C-terminus is sufficient to bind AMBRA1^WD40^, we performed a series of pull-down assays. Cyclin D1 residues P268-I295 were directly fused to GST and co-expressed with CDK4_EE_-His, TSF-DDB1 and MBP-AMBRA1^WD40^. Using GST-cyclin D1 (residues P268-I295) construct as bait was sufficient to pull-down AMBRA1^WD40^. The same result was obtained by using MBP-AMBRA1^WD40^ as a bait, showing that the cyclin D1 C-terminal sequence is sufficient for recognition by AMBRA1^WD40^. The western blotting analysis using anti-p-cyclin D1 antibody confirmed presence of T286 phosphorylation in all pulled samples (Fig. 4A). We therefore tested the requirement for T286 phosphorylation using T286A mutants. T286A mutation largely decreased the interaction with AMBRA1^WD40^, indicating phosphorylation-dependent interaction (Fig. 4A). Subsequently, we tested whether cyclin D1 C-terminus is necessary for the interaction. For this, we co-transfected the three truncated TSF-cyclin D1 constructs (residues M1-D267, M1-A271 or M1-D282) individually with GST-CDK4_EE_, His-DDB1 and MBP-AMBRA1^WD40^. The C-terminal truncations almost completely abolished the interaction with AMBRA1^WD40^, indicating that the cyclin D1 C-terminus is as well necessary for the interaction with AMBRA1^WD40^ (Fig. 4B). Next, we performed pull-down assays using full-length cyclin D1, cyclin D2 and cyclin D3. All three D-type cyclins were able to associate with AMBRA1^WD40^, while the corresponding threonine phosphorylation site mutations (cyclin D1 T286A, cyclin D2 T280A and cyclin D3 T283A) almost completely abolished AMBRA1 binding (Fig. 4C). Therefore, the C-terminal phosphodegron of D-type cyclins is the key region responsible for interaction with the AMBRA1 WD40 domain. To examine whether AMBRA1^WD40^ can mediate ubiquitination of D-type cyclins, we performed *in vitro* ubiquitination assays using TSF-D-type cyclins, and the corresponding threonine phosphorylation site mutants. Consistent with the pull-down results, all three D-type cyclins were poly-ubiquitinated when incubated with the CRL4 ligase complex (Fig. 4D, E), and the threonine phosphorylation site mutants showed reduced ubiquitination activity in all three D-type cyclins (Fig. 4D, E). Furthermore, the full-length AMBRA1 was also used to perform the *in vitro* ubiquitination assay. The results showed the similar trends as observed for the AMBRA1^WD40^ (Fig. 4F), indicated that the AMBRA1 WD40 domain is sufficient to mediate ubiquitination of D-type cyclins.

**Fig. 4.**
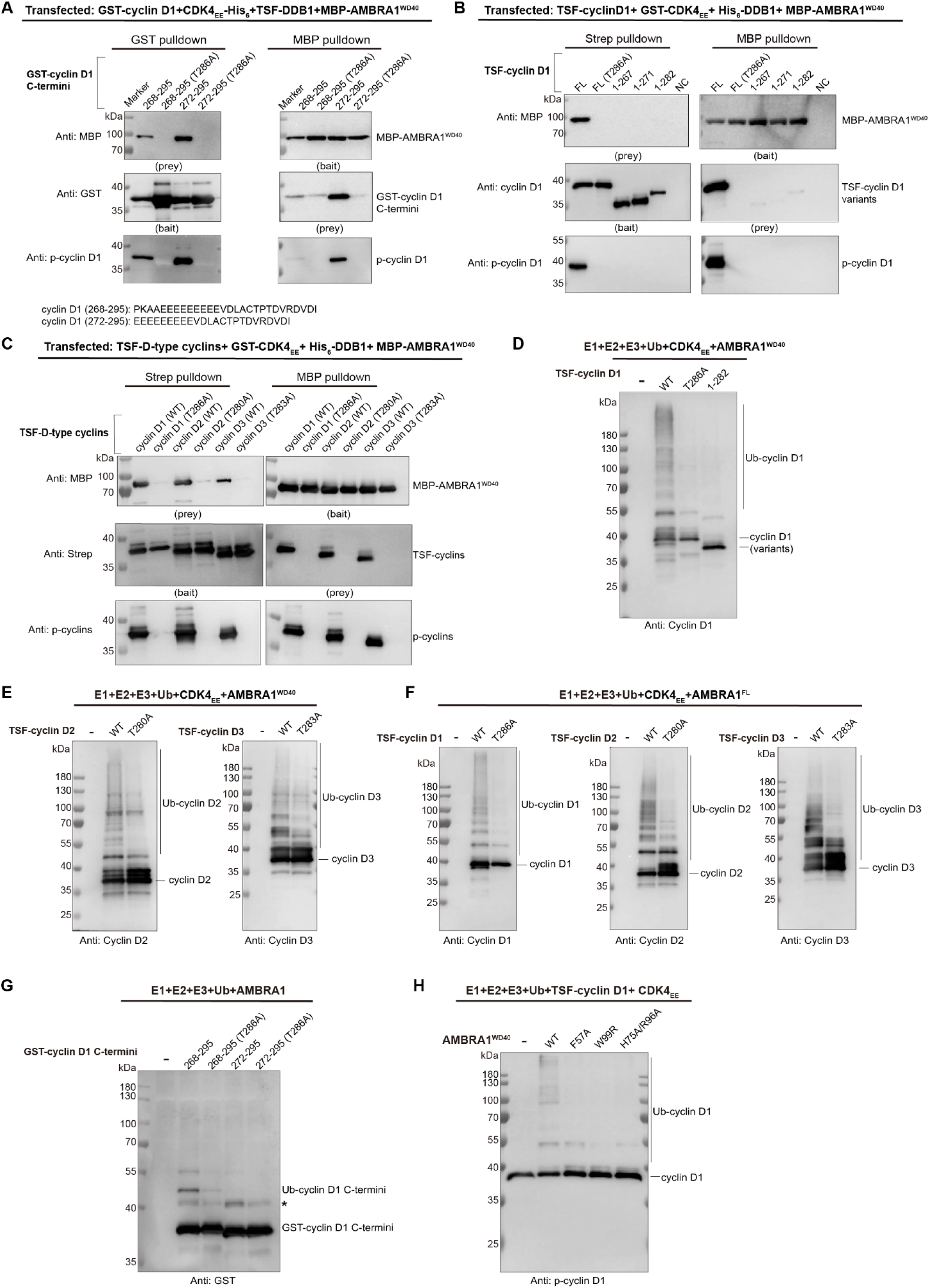
AMBRA1 ^WD40^ mediates ubiquitination of D-type cyclins. **(A)** *In vitro* pull-down experiment of MBP-AMBRA1^WD40^ with GST tagged C-terminal region of cyclin D1 and T286A mutants. The experiment was repeated at least three times and visualized by western blotting. **(B)** *In vitro* pull-down experiment of MBP-AMBRA1 ^WD40^ with cyclin D1 C-terminal truncations and T286A mutants. The cells were co-transfected with TSF-cyclin D1, GST-CDK4_EE_, His_6_-DDB1 and MBP-AMBRA1^WD40^.The experiment was repeated at least three times and visualized by western blotting. **(C)** *In vitro* pull-down experiment of MBP-AMBRA1^WD40^ with cyclin D1, cyclin D2, cyclin D3 and cyclin D1^T286A^, cyclin D2^T280A^ and cyclin D3^T283A^ mutants. The cells were co-transfected with TSF-D-type cyclins, GST-CDK4_EE_, His_6_-DDB1 and MBP-AMBRA1^WD40^.The experiment was repeated at least three times and visualized by western blotting. **(D)** Ubiquitination assay of AMBRA1^WD40^ mediated the polyubiquitination of TSF-cyclin D1, T286A mutant and C-terminal truncation. The experiment was repeated independently three times with similar results. **(E)** Ubiquitination assay of AMBRA1^WD40^ mediated the polyubiquitination of TSF-cyclin D2/ cyclin D2^T280A^ and TSF-cyclin D3/ cyclin D3^T283A^. The experiment was repeated independently three times with similar results. **(F)** Ubiquitination assay of AMBRA1^FL^ mediated the polyubiquitination of TSF-D-type cyclins and threonine phosphorylation site mutants. The experiment was repeated independently three times with similar results. **(G)** Ubiquitination assay of AMBRA1 mediated the polyubiquitination of GST-cyclin D1 C- termini. Asterisk indicates the non-specific band. The experiment was repeated independently three times with similar results. **(H)** Ubiquitination assay of AMBRA1^WD40^ WT and key binding sites mutants mediated the polyubiquitination of cyclin D1. The results were analysed by western blotting.

Previous studies have shown that residue K269, which is located in the proximity of the C-terminal phosphodegron, is essential for the ubiquitination of cyclin D1 by the SCF ligase, resulting in its degradation (*24*). Similarly, more recent post-translational modification analysis suggests that the T286 plays an important regulatory role in controlling the ubiquitination of cyclin D1 at the K269, as the major site for this modification (*7*). To address whether AMBRA1^WD40^ can mediate ubiquitination of the cyclin D1 C-terminus at K269, we performed *in vitro* ubiquitination assays using GST-cyclin D1 (residues P268-I295) and GST-cyclin D1 (residues E272-I295) constructs. Both constructs were able to associate with AMBRA1^WD40^ (Fig. 4A), however only the construct that includes K269 (GST-cyclin D1 (residues P268-I295)) showed robust ubiquitination when incubated with the CRL4 ligase complex. Consistent with the previous studies, the corresponding T286A mutant showed strongly reduced ubiquitination activity (Fig. 4G).

To address whether AMBRA1^WD40^ F57A, W99R and H75A/R96A mutants influence the substrate ubiquitination, we purified the AMBRA1^WD40^ mutants in complex with Cullin4A-DDB1-RBX1 E3 ligase, and CDK4_EE_-cyclin D1 complex and performed *in vitro* ubiquitination assays (Fig. S1B, C). AMBRA1^WD40^ is sufficient to promote ubiquitination of cyclin D1 if every enzyme of the ubiquitination cascade reaction is included (Fig. 4D, H). However, the polyubiquitination of cyclin D1 was strongly decreased in the presence of all tested AMBRA1^WD40^ mutants when compared with the wild-type protein, which is consistent with the pull-down results (Fig. 2D, 4H). Taken together, our results imply that AMBRA1^WD40^ is sufficient for phosphorylation-dependent association with the cyclin D1 C-terminus and subsequent ubiquitination at K269.

### Mutations that disrupt the AMBRA1-cyclin D1 interaction cause DNA damage

We further examined the effect of the interaction between cyclin D1 and AMBRA1 on the DNA damage in U2OS cells. It was reported that AMBRA1 depletion leads to an increase in cellular proliferation, the S-phase enrichment, replication stress and DNA damage, which is greatly intensified by inhibition of CHK1 (*8*). Short interfering RNA (siRNA) oligoribonucleotides were used to knockdown endogenous AMBRA1 in U2OS cells (Fig. 5A). Consistent with previous studies, we found that AMBRA1 downregulation results in the higher expression levels of cyclin D1 and also significantly increased DNA damage in cells treated with CHK1 inhibitor AZD7762 (Fig. 5A, B). We have investigated the effect of over-expression the wild-type and mutated AMBRA1 on endogenous DNA damage in AMBRA1-silenced cells. Re-expression of wild-type AMBRA1 was sufficient to rescue the non-interfered cell phenotype and reduce the cyclin D1 expression level (Fig. 5A-C). All tested AMBRA1 mutants failed to attenuate the high levels of endogenous DNA damage and downregulate the cyclin D1 expression level (Fig. 5A-C). In summary, F57, W99, H75 and R96 are the key binding sites, which are essential for AMBRA1- mediated ubiquitination and degradation of cyclin D1, thereby directly impacting the essential downstream cellular functions. These results highlight the critical role of AMBRA1 in regulation of cyclin D1 protein level in cells, and demonstrate the importance of the AMBRA1 in preventing replication stress and genome instability.

**Fig. 5.**
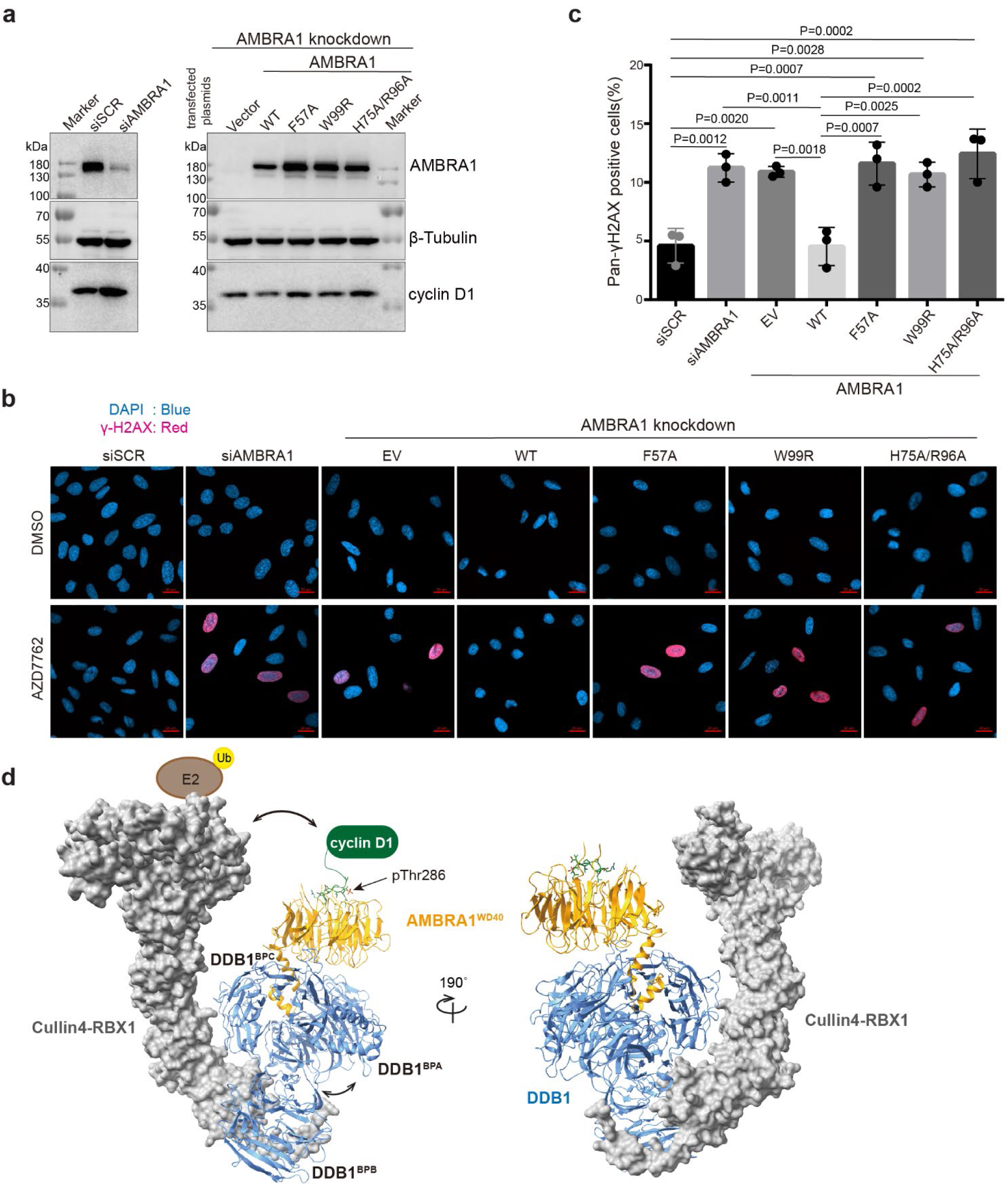
AMBRA1 mutants result in increased sensitivity to CHK1 inhibition. **(A)** AMBRA1 was knocked down by specific short interfering RNA (siRNA) in U2OS cells. Then cells were transfected with the empty vector, AMBRA1 WT and mutants. The protein expression level was analysed by western blotting, β-tubulin was used as the protein loading control. siSCR, scrambled siRNA. WT, wild-type. **(B)** The representative images of Pan-γH2AX-positive cells after AMBRA1 knockdown, AMBRA1 WT and mutants overexpression. Cells were treated with DMSO or 100 nM AZD7762. Imaging was collected using a Zeiss LSM 900 confocal microscope. DAPI was shown in blue, γH2AX was shown in red. Scale bar = 20 μm. **(C)** Quantification of Pan-γH2AX-positive cells. For each group, around 300 cells were manually counted by using ZEN software, the percentage of pan-γH2AX positive cells were calculated. Data was plotted in GraphPad Prism 9.4.1 software. Statistical significance was determined by Tukey’s multiple comparisons of one-way ANOVA, dots indicate individual data points for n = 3 biological replicates. P values are indicated in the graph. Graph bar was presented as mean values +/− SD. **(D)** Hypothetic model of Cullin4-RBX1-DDB1-AMBRA1^WD40^-cyclin D1 complex. The AMBRA1^WD40^-cyclin D1 structure in this study aligned to Cullin4A-DDB1-RBX1 structures (PDB: 2HYE). Cullin4 and RBX1 are presented on the surface, and DDB1, AMBRA1 and cyclin D1 are colored in blue, orange and green, respectively. The phosphorylated Thr286 is indicated with the arrow. The movement of DDB1 between BPA-BPC with BPB facilitates the E2 close to substrate that allows the ubiquitination of cyclin D1.

## Discussion

The overexpression and amplification of D-type cyclins is frequently found in several types of human cancer (*1*). Emerging evidence indicates that AMBRA1 is a major regulator of the stability of D-type cyclins, including cyclin D1, D2, and D3, through interaction with DDB1 and CRL4 (*6–8*). By promoting their degradation, AMBRA1 helps maintain proper cyclin D levels, preventing overaccumulation that can lead to uncontrolled cell proliferation. Understanding their interaction may provide critical insights into therapeutic targets for cancers with altered cell cycle regulation. Our previous studies have shown that AMBRA1 interacts with the DDB1 double propeller via multiple interfaces, which involve the N-terminal helix-loop-helix motif and the bottom surface of the AMBRA1 WD40 domain (*15*). This bipartite interaction mechanism is accompanied by the stabilization effect on the AMBRA1 “split” WD40 domain dynamics. In this study, we determined the structure of the DDB1-AMBRA1-cyclin D1 complex, and identified the WD40 domain as a crucial structural element used by AMBRA1 to recognize the D-type cyclins. AMBRA1 interactions with most of the binding partners, including Beclin1, dynein light chain 1 (DLC1), LC3 and PP2A, are mediated by sequences that localize to the intrinsically disorder regions (*14, 19, 25–27*). Interaction with DDB1 is proposed to stabilize the split WD40 domain in AMBRA1, and bring these regions to the proximity of the CRL4 active site. Our study now shows that AMBRA^WD40^ can directly recognize and bind to a highly conserved extreme C-terminus of cyclin D1. Cyclin D1 C-terminus folds into a turn-like conformation and tightly packs against the top surface of the AMBRA1 β-propeller, indicating a general mode for recognition of D-type cyclins. Our results indicate that the complete, fully-formed β-propeller assembly of the AMBRA1 WD40 domain is necessary for it to recognize and mediate ubiquitination of D-type cyclins, and interaction with DDB1 may be an important stabilizing factor.

D-type cyclins, share about 57% sequence identity, and contain a conserved C-terminal phosphodegron that determine their degradation by the UPS (*1*). Using *in silico* predictions and biochemical assays, we found that AMBRA^WD40^ is likely able to recognize and bind to cyclin D2 and cyclin D3 in a manner similar to how it binds to cyclin D1. The structure shows that cyclin D1 residue T286 is phosphorylated in the reconstituted complex and plays a pivotal role in cyclin D1 interaction with the AMBRA1 WD40 domain. Cyclin D1 (T286A) mutation prevents interaction with AMBRA1^WD40^ and impair ubiquitination near the C-terminus of cyclin D1. Our findings agree with previous studies demonstrating that in contrast to the wild-type cyclin D1, depletion of AMBRA1 does not induce accumulation of cyclin D1 (T286A) (*7*). In addition to this key residue, AMBRA1 F57 and W99 residues are lining the hydrophobic ligand binding pocket, which, combined with the T286-interacting residues H75 and R96 complete the cyclin D1 recognition site. Mutations of any of these residues impaired AMBRA1^WD40^ ability to interact and mediate ubiquitination of cyclin D1. Furthermore, none of the mutants was able to rescue the increased DNA damage in the AMBRA1 silenced cells, consistent with cyclin D1 stabilization. These observations underscore the importance of AMBRA1^WD40^-cyclin D1 interaction in regulating cell proliferation, by mediating cyclin D1 degradation, and prevention of replication stress and DNA damage. The general mode of cyclin D1 recognition is consistent with previous studies showing that the top surface of WD40 domains frequently interact with proteins that are targeted for ubiquitination (reviewed in (*28*)). We propose a model in which phosphorylation on Thr286 that activates cyclin D1 nuclear export (*10*) is required for its association with AMBRA1^WD40^. This interaction mode results in cyclin D1 presentation toward the RBX1 binding site of the E2 ubiquitin-conjugating enzyme (Fig. 5D). Neddylation of Cullin likely induces RBX1 to adopt an extended catalytic conformation (*29*), thereby facilitating the transfer of ubiquitin from the E2 enzyme to cyclin D1.

Our structural and biochemical findings uncovered a detailed mechanism of CRL4^AMBRA1^-mediated degradation of D-type cyclins. The CRL4^AMBRA1^ complex plays a crucial role in targeting D-type cyclins for ubiquitination and subsequent proteasomal degradation. By understanding the molecular interactions and conformational changes that occur during this process, our study sheds light on the broader implications of CRL4^AMBRA1^ activity in cell cycle control. Future studies are needed to determine whether AMBRA1^WD40^ can directly recognize substrates other than cyclins D and functional role of the intrinsically disordered regions. Small molecular compounds like molecular glues and PROTACs (Proteolysis Targeting Chimeras) are at the forefront of targeted protein degradation technology. They function by promoting or enhancing the interaction between E3 ubiquitin ligases and target proteins, leading to the ubiquitination and proteasomal degradation of the substrate. Structural studies of the AMBRA1 and D-type cyclins have laid the foundation for the development of small molecules targeting the E3 ligase-receptor-substrate complex, and provide the potential treatment for diseases characterized by dysregulated cyclin activity, such as cancer.

## Materials and Methods

### Antibodies

Anti-AMBRA1 antibody was used at 1:1000 dilution (CST, 24907s), anti-Phospho-Histone H2A.X (Ser139) antibody was used at 1:1000 dilution (CST, 2577s), Fab2 Fragment (Alexa Fluor 647 Conjugate) was used at 1:1000 dilution (CST, 4414s), anti-cyclin D1 antibody was used at 1:1000 dilution (CST, 2922S and Santa Cruz, sc-20044), anti-cyclin D2 antibody was used at 1:1000 dilution (Proteintech, 10934-1-AP), anti-cyclin D3 antibody was used at 1:1000 dilution (Proteintech, 26755-1-AP), anti-Phospho-cyclin D1 (Thr286) antibody was used at 1:2000 dilution (CST, 3300T), anti-Phospho-cyclin D2 (Thr280) antibody was used at 1:2000 dilution (Abcam, AB230818), anti-Phospho-cyclin D3 (Thr283) antibody was used at 1:2000 dilution (Affinity, AF3251), anti-β-Tubulin antibody was used at 1:1000 dilution (Novus, NB600-936), anti-MBP tag Mouse Monoclonal antibody was used at 1:1000 dilution (GenScript, A00190), GST tag Mouse Monoclonal antibody was used at 1:1000 dilution (STARTER, S-372-19), the secondary antibodies conjugated to horseradish peroxidase (HRP), Goat anti-Mouse IgG was used at 1:2000 dilution (CWBIO, CW0102), Goat anti-Rabbit IgG was used at 1:2000 dilution (CWBIO, CW0103).

### Cloning and mutagenesis

The gene for full-length AMBRA1 (residues M1-R1298) was codon optimized. The genes encoding DDB1, Cullin4A, RBX1, cyclin D1, CDK4, UBA3, APPBP1, UBC12 and NEDD8 were amplified from cDNA by PCR and subcloned into pCAG vectors with different tags using KpnI and XhoI cutting sites, respectively. AMBRA1^WD40^ (residues M1-N204/S853-G1044) was subcloned with N- terminal MBP tag or GST tag followed by a TEV cutting site. For cyclin D1, cyclin D2 and cyclin D3, all were cloned with N-terminal Twin-Strep-FLAG (TSF) tag. DDB1 was constructed with N- terminal His_6_ tag, or N-terminal TSF tag. Residues G43-G47 in CDK4 were replaced with Glu-Glu (CDK4_EE_) in order to increase the protein stability (*20*). CDK4_EE_ as constructed with N-terminal GST tag followed by a TEV cutting site. Cullin4A and RBX1 were constructed with N-terminal TSF tag.

Wild-type, truncations and mutations of cyclin D1 were constructed with N-terminal TSF tag for strep pull-down. The C-terminal region was cloned with N-terminal GST tag and followed by a (GGGGS)_2_ linker used for GST pull-down.

The AMBRA1^WD40^ mutants were generated by PCR-based site-directed mutagenesis approaches. For AMBRA1^WD40^ WT, AMBRA1^WD40^ F57A, F57R, W99A, W99R, H75A/R96A, F57A/W99A, F57R/W99A, F57A/W99R, H75A/R96A/W99A and H75A/R96A/W99R used in the MBP pull-down experiment, they were cloned as a N-terminal MBP tag. For co-expression with Cullin4A- DDB1-RBX1 E3 ligase, AMBRA1^WD40^ WT, AMBRA1^WD40^ F57A, W99R and H75A/R96A was constructed with N-terminal GST tag. For *in vivo* cell experiments, tagless AMBRA1^FL^, AMBRA1^F57A^, AMBRA1^W99R^ and AMBRA1^H75A/R96A^ were cloned into pCAG vector. The primers used in this study are listed in Table S1.

### Protein expression and purification

For mammalian cell expression, MBP-AMBRA1^WD40^, His_6_-DDB1, TSF-cyclin D1 and GST- CDK4_EE_ were co-transfected and purified for structural studies. GST-AMBRA1^WD40^ (WT and mutants), TSF-Cullin4A, TSF-DDB1 and TSF-RBX1 were co-transfected and purified for *in vitro* ubiquitination assay. TSF-cyclin D1 (WT and mutants) and GST-CDK4_EE_ were co-expressed for *in vitro* ubiquitination assay. MBP-AMBRA1^WD40^ (WT and mutants) and His_6_-DDB1 were co-expressed for negative staining.

The Expi293F cells were grown in Union-293 medium and used for protein expression, with 1mg DNA and 4mg PEI per 1 liter of cells at density of 2 × 10^6^ cells/ml. Cells were harvested after transfected for 3 days and washed with 1× cold PBS.

For AMBRA1^WD40^, DDB1, cyclin D1 and CDK4_EE_ co-expression, the cell pellet was lysed in lysis buffer A (50 mM HEPES pH 7.4, 200 mM NaCl, 10% glycerol (v/v), 1% Triton-X100 (v/v), 2 mM MgCl_2_, 5 mM beta-mercaptoethanol) with proteases inhibitors (1 mM PMSF, 0.15 μM aprotinin, 10 μM leupeptin, 1 μM pepstain) for 30min at 4 °C. After centrifugation at 39,191 × g for 30 min, MBP beads were incubated with the cell supernatant at 4 °C for 2 h. The beads were washed 20 bed volumes with wash buffer B (50 mM HEPES pH 7.4, 200 mM NaCl, 2 mM MgCl_2_, 5 mM beta-mercaptoethanol). Bound proteins were eluted with wash buffer B containing 10 mM maltose. After TEV digestion overnight, the elution was then subjected to Strep-tactin sepharose resin, washed 20 bed volumes with wash buffer B and eluted with gel filtration buffer C (50 mM HEPES pH7.4, 200 mM NaCl, 2 mM MgCl_2_ and 1 mM TCEP) supplemented with 5 mM desthiobiotin. The protein elution was further purified by Superdex 200 Increase 10/300 GL (Cytiva). The sample was used for cryo-EM structure analysis. The SDS-PAGE for the purified proteins was shown in Fig. 1C and the gel filtration profile was shown in Fig. S1A.

The AMBRA1^WD40^ (WT and mutants) and Cullin4A-DDB1-RBX1 E3 ligase complex was purified as previously described (*15*). In brief, the cell pellet was lysed in lysis buffer A with proteases inhibitors (1 mM PMSF, 0.15 μM aprotinin, 10 μM leupeptin, 1 μM pepstain) for 30 min at 4 °C. After centrifugation at 39,191 × g for 30 min, GST beads were incubated with the cell supernatant at 4 °C for 2 h. The beads were washed 20 bed volumes with wash buffer B and eluted in wash buffer B containing 50 mM reduced glutathione. After TEV digestion overnight, the elution was further applied to Strep-tactin sepharose resin, washed 20 bed volumes and eluted with gel filtration buffer C containing 5 mM desthiobiotin. The purified proteins are shown in Fig. S1B.

For cyclin D1/ cyclin D2/ cyclin D3 (WT and mutants)-CDK4_EE_ complex, the cell pellet was lysed in lysis buffer A with proteases inhibitors (1 mM PMSF, 0.15 μM aprotinin, 10 μM leupeptin, 1 μM pepstain) for 30 min at 4 °C. After centrifugation at 39,191 × g for 30 min, GST beads were incubated with the cell supernatant at 4 °C for 2 h. The beads were washed 20 bed volumes with wash buffer B and eluted in wash buffer B containing 50 mM reduced glutathione. After TEV digestion overnight, the elution was further applied to Strep-tactin sepharose resin, washed 20 bed volumes and eluted with gel filtration buffer C containing 5 mM desthiobiotin. The SDS-PAGE for the purified proteins was shown in Fig. S1C, G. GST-linker-cyclin D1 C-termini proteins were purified by one-step GST affinity purification (Fig. S1F). The eluted proteins were further used for *in vitro* ubiquitination activity assay.

For AMBRA1^WD40^-DDB1 complex, the cell pellet was lysed in lysis buffer A with proteases inhibitors for 30 min at 4 °C. After centrifugation at 39,191 × g for 30 min, MBP beads were incubated with the cell supernatant at 4 °C for 2 h. The beads were washed 20 bed volumes with wash buffer B and eluted in wash buffer B containing 10 mM maltose. After TEV digestion overnight, the elution was concentrated and injected into Superdex 200 Increase 10/300 GL (Cytiva) (Fig. S1D, E). The purified proteins were further used for negative staining.

### Negative stain EM imaging

Four microliters of purified protein (0.03 mg/ml) was loaded onto a freshly glow-discharged carbon-coated 300 mesh grid (EMCN, BZ11023a) for 1 minute. The grids were subsequently blotted with filter paper and negatively stained with 2% (w/v) uranyl acetate for 40 seconds. Redundant liquid was absorbed using filter paper. The sample was imaged using a Talos 120C transmission electron microscope (Thermo Fisher Scientific) performed at 120 kV in low-dose mode and imaged with a Ceta CMOS camera (Thermo Fisher Scientific). Data were collected at a nominal magnification of 73,000X, corresponding to a pixel size of 1.96 Å. The micrographs were further processed by cryoSPARC4.4.1 (*30*).

### Cryo-EM grid preparation and data acquisition

Human DDB1-AMBRA1^WD40^-cyclin D1-CDK4_EE_ complex was prepared for cryo-EM data acquisition. The protein complex was eluted from the peak of size exclusion column. For cryo-EM grid, 3 μL of 0.3 mg/ml protein sample were deposited onto freshly glow-discharged Quantifoil R1.2/1.3 Cu300 mesh grids and plunged into liquid ethane using a FEI Vitrobot Mark IV after blotting for 3 s with blot force 0 at 4 °C and 100% humidity. A total of 6,120 movies were collected on a Titan Krios electron microscope operating at 300 kV equipped with Gantan K3 camera at a defocus of −1.0 μm to −1.8 μm in counting mode, corresponding to a pixel size of 0.85 Å. Automated image acquisition was performed using SerialEM with a 3 × 3 image shift pattern. Movies consists of 50 frames, with a total dose of 59.9 e^−^/ Å^2^, a total exposure time of 2.5 s and a dose rate of 17.32 e^−^/pixel/sec. Imaging parameters for the dataset are summarized in Table S2.

### Cryo-EM data processing

The movies were divided into five-by-five patches for motion correction by Relion’s own implementation of the MotionCor2 algorithm, and then imported into cryoSPARC v4.4.1 (*30–33*). CTF fitting and estimation were performed by patch CTF estimation. After particle picking from an initial subset of 500 micrographs with blob picker, 322,822 particles were extracted and subjected to iterative rounds of 2D classifications. About 3.7 millions of particles were picked from the remaining 5,620 micrographs. All the picked particles were extracted with a box size of 256 x 256 pixels and further binned to 128 x 128 pixels to speed up the data processing. After iterative 2D classifications, the cleaned 2,794,307 particles were further subjected for *ab initio* to generate four different classes and heterogeneous refinement. The particles from the best three classes were re-extracted with a box size of 256 x 256 pixels and chosen for homogeneous refinement. Due to the flexibility of the BPB domain of DDB1, we performed particle subtraction to remove the signal of this domain from particle images, followed by local refinement. After cycles of 2D classifications and one more round of heterogeneous refinement, the best resolved class was chosen for non-uniform refinement. The final map obtained from the 1,015,935 particles was estimated at 3.55 Å. The map was post-processed by deepEMhancer with tight target modes and used for model building (*34*). All reported resolutions are based on the gold-standard FSC 0.143 criterion.

### Atomic model building and refinement

The coordinates for DDB1-AMBRA1^WD40^ (8WQR) were rigid body fitted into the deepEMhancer post-processed map using UCSF Chimera and the residues of cyclin D1 were manually built using Coot (*35*). Atomic coordinates were refined by iteratively performing Phenix real-space refinement using the unsharpened map (*36*). Manual inspection and correction of the refined coordinates were performed in Coot (*35*). Model quality was assessed using MolProbity and the map-vs-model FSC by comparing the map-vs-model FSC with the FSC of the experimental cryo-EM density (Fig. S5). Figures were prepared using PyMOL2.3.3 (The PyMOL Molecular Graphics System, Version 2.3.3 Schrödinger, LLC), UCSF Chimera version 1.15 and ChimeraX version 1.3 (*37*). The cryo-EM density map has been deposited in the Electron Microscopy Data Bank under accession code EMD- 60925 and the coordinate has been deposited in the Protein Data Bank under accession number 9IVD.

### Neddylation of Cullin4A with HA tagged NEDD8

Neddylation of Cullin4A was performed by co-expressing CRL4^AMBRA1^ ^FL/WD40^ with tagless APPBP1, UBA3, UBC12 and HA tagged NEDD8. For CRL4^AMBRA1^ ^FL^ neddylation, MBP- AMBRA1-His_6_ was co-transfected with Cullin4A-DDB1-RBX1 E3 ligase complex, tagless APPBP1, UBA3, UBC12 and HA tagged NEDD8, then purified as previously described AMBRA1^FL^-DDB1-Cullin4A-RBX1 complex purification (*15*). For CRL4^AMBRA1^ ^WD40^ neddylation, GST-AMBRA1^WD40^ was co-transfected with Cullin4A-DDB1-RBX1 E3 ligase complex, tagless APPBP1, UBA3, UBC12 and HA tagged NEDD8, then purified as previously described AMBRA1^WD40^-DDB1-Cullin4A-RBX1 complex purification (*15*). Neddylation detection was performed by western blotting; the purified proteins were further used for Cyclin D *in vitro* ubiquitination assay (Fig. 4E, G, H). The SDS-PAGE and western blotting for the purified proteins were shown in Fig. S6.

### *In vitro* ubiquitination assay

The ubiquitination assays were performed in a 25 μL reaction volume with the following components: 100 nM UBE1 (Boston Biochem E-304), 1.5 μM UBCH5C (Boston Biochem E2- 627), 0.3 μM purified AMBRA1^WD40^-DDB1-Cullin4-RBX1 or AMBRA1^FL^-DDB1-Cullin4-RBX1 complex, 20 μM HA-ubiquitin (Boston Biochem U-110), 0.16 μM TSF-D-type cyclins-CDK4_EE_ complex or GST-cyclin D1 C-termini, and 10 mM MgATP solution (Boston Biochem B-20) in E3 ligase reaction buffer (Boston Biochem B-71). The reaction was incubated at 37 °C for 2 h and analysed by SDS-PAGE, followed by immunoblot. The experiment was repeated three times with similar result.

### RNA interference

siRNA oligoribonucleotides corresponding to the human AMBRA1 were ordered from Tsingke. AMBRA1 siRNA 1: 5′- GAGUAGAACUGCCGGAUAG−3′; AMBRA1 siRNA 2: 5′- CCACCCAUGUGAACCAUAA-3′; AMBRA1 siRNA 3: 5′-GCGGAGACAUGUCAGUAUC-3′; AMBRA1 siRNA4: 5′-CUGAAUCGCUGUCGUGCUU-3′; For siRNA transfection, the human osteosarcoma U2OS cells were plated at 7 × 10^5^ cells per well in 6-well plate. Each well transfected with optimal amount of siRNA mixture by using Lipofectamine RNAi-MAX (ThermoFisher Scientific) next day, according to the manufacturer’s instructions.

### Immunoblotting and immunofluorescence

The U2OS cells were plated into 6-well plate, then followed by siRNA transfection using Lipofectamine RNAiMAX (ThermoFisher Scientific, 13778075) next day, according to the manufacturer’s instructions. After siRNA transfection for 24 h, cells were plated into 12-well plate for transient plasmids transfections next day. The transient plasmids transfections were performed using X-tremeGENE HP DNA Transfection Reagent (Roche, 06366236001) according to the manufacturer’s instructions. Cells were harvested 48 h after plasmids transfection. Cells were lysed in lysis buffer (50 mM Tris pH 7.4, 150 mM NaCl, 0.5 mM EDTA, 1% NP-40, 0.1% SDS) supplemented with protease inhibitors (1 mM PMSF, 0.15 μM aprotinin, 10 μM leupeptin, 1 μM pepstain) for 15 min on ice, then centrifuged at 15,000 × g for 15 min to remove insoluble debris. Solubilized proteins were quantified by BCA Protein Assay Kit (Sangon Biotech, C503021), equal amounts of protein were mixed with loading buffer and heated at 95 °C for 5min. Samples were loaded at same amount by SDS-PAGE and transferred to 0.22 μm Immuno-Blot PVDF membranes (BIORAD). Membranes were then blocked in 5% milk/TBST for 1 h at room temperature and incubated with primary antibodies at 4 °C overnight. For the detection of proteins, using the secondary antibodies conjugated to horseradish peroxidase (anti-mouse and antirabbit, CWBIO) at 1:2000 dilution in TBST for 1 h at room temperature and visualizing with cECL Western Blotting Kit (CWBIO, CW0048). The images were acquired by Amersham Imager 680.

For immunofluorescence, U2OS cells were plated into the 12-well plate on coverslips at 5 × 10^4^ density per well, then followed the same procedure as for immunoblotting. For CHK1 inhibition, cells were treated with 100 nM of AZD7762 (MedChemExpress, HY-10992) for 24 h after plasmids transfection. Cells were washed with cold PBS and fixed in 4% Paraformaldehyde (PFA, Sangon, E672002) in PBS for 20 min at room temperature, then permeabilized with PBS/0.1% (v/v) Triton X-100 for 20 min and subjected to blocking with PBS/1% (w/v) BSA to block the cell for 1 h at room temperature. γH2AX antibody were diluted in PBS/0.01% (v/v) Triton X-100 at 1:1000 dilution, applied to the coverslips at 4 °C overnight. Alexa647-conjugated secondary antibody was diluted in PBS/1% (w/v) BSA at 1:1000 dilution and incubated with the coverslips for 1 h at room temperature. Slides were mounted in Anti-Fade Mounting Medium with DAPI (Beyotime, P0131). Imaging was performed using a Zeiss LSM 900 confocal microscope with a Plan-Apochromat 10×/0.95 and 40×/0.95 objective.

### *In vitro* pull-down and western blotting experiment

MBP-AMBRA1^WD40^ (WT and mutants), His_6_-DDB1, TSF-cyclin D1/ TSF-cyclin D2/ TSF-cyclin D3 and GST-CDK4_EE_ were co-transfected into Expi293F cells. The cells were harvested after 3 days of transfection and washed with 1 x PBS. After lysis and centrifugation, the supernatant was divided into halves and incubated with MBP or Strep-tactin sepharose resin for 2 h. The beads were washed with lysis buffer A for 2 times, and with wash buffer B for 2 times, further eluted with gel filtration buffer C with 10 mM maltose or 5 mM desthiobiotin. The eluted sample were detected by SDS-PAGE and wester blotting with antibodies against MBP tag and cyclin D1.

To test the C-terminal region of cyclin D1 interact with AMBRA1^WD40^, GST-linker-cyclin D1 (C-termini), MBP-AMBRA1^WD40^, TSF-DDB1 and CDK4_EE_-His_6_ were co-transfected into Expi293F cells. The GST and Strep pull-down assay and western blotting were performed using the same protocol.

To test the C-terminal truncation of cyclin D1 interact with AMBRA1^WD40^, TSF-cyclin D1 (WT, T286A, 1-282, 1-271 and 1-267), MBP-AMBRA1^WD40^, His_6_-DDB1, and GST-CDK4_EE_ were co-transfected into Expi293F cells. The MBP and Strep pull-down assay and western blotting were performed using the same protocol.

The experiment was repeated three times independently.

### Identification of phosphorylation sites by mass spectrometry

The purified protein complex was digested on an immobilized pepsin column and eluted peptides were desalted using a trap column (1mm x 15mm, Acclaim PepMap300 C18, 5 μm, Thermo Fisher Scientific) for 5 minutes at a flowrate of 200 μl/min, with a mobile phase consisting of 0.1% formic acid. Subsequent peptide separation was performed on the ACQUITY BEH C18 (2.1 x 50 mm) analytical column with a first gradient ranging from 9 to 45% of buffer B (80% acetonitrile and 0.1% formic acid) for 10 minutes followed by a second gradient ranging from 45 to 99% of buffer B for 1 minute, at a flowrate of 50 μl/min. Peptides were directly ionized via electrospray ionization and analysed by Orbitrap Eclipse (Thermo Fisher) mass spectrometer. A full MS scan was collected in the Orbitrap (m/z: 375–1500, resolution: 60,000, AGC 4e5, max injection time: 118 ms) followed by a series of MS/MS scans measured in the Orbitrap (HCD normalized collision energy: 30%, resolution: 30,000, AGC 5e4, isolation width 1.6 m/z, max injection time: 54ms). Unknown and singly charged ions were excluded from fragmentation. All MS/MS spectra were searched against the custom database containing the sequences of the DDB1, AMBRA1^WD40^, cyclin D1 and CDK4 and common contaminant proteins (CRAPome/Strep tag (*38*)) using the Proteome Discoverer 2.5 software (Thermo Fisher). The following parameters were used: MS tolerance, 10 ppm; MS2 tolerance, 0.02 Da; missed cleavages, 2; No-Enzyme (Unspecific); variable modifications, oxidation methionine (+15.995 Da), Phosphorylation (S, T, Y, +79.966), and N-terminal acetylation (+42.011 Da). The Percolator node within Proteome Discoverer was used to filter the peptide spectral match (PSM) false discovery rate to 1 % (strict). The ptmRS node was used for localization of modification sites within validated peptide sequences (*39*).

### Quantification and statistical analyses

For Pan-γH2AX positive cells quantification, a minimum of three independent experiments were included in the representing graphs. For each group, total around 300 cells were manually counted by using ZEN (Blue edition) software, pan-γH2AX positive cells percentage were calculated. Data were analysed using GraphPad Prism 9.4.1 software. Statistical significance was determined by Tukey’s multiple comparisons of one-way ANOVA, P values are indicated in the graph, the value of p < 0.05 was considered statistically significant. Graph bar was presented as mean values +/− SD.

## Acknowledgments

The authors thank the cryo-EM center (KEMC) and advanced mass spectrometry facility (KMS) of Kobilka Institute of Innovative Drug Discovery, the Chinese University of Hong Kong (Shenzhen) for the support.

## Author contributions

Conceptualization: YW, ML, MS, GS

Methodology: YW, ML, MS, GS

Investigation: YW, ML, SW, FT, XW, XM, TL, MS, GS

Visualization: YW, ML, XM, MS, GS

Supervision: MS, GS

Writing—original draft: MS, GS

Writing—review & editing: YW, ML, FT, XW, XM, MS, GS

## Data and materials availability

The atomic coordinates of the AMRBA1^WD40^-DDB1 bound to cyclin D1 reported in this study and associated cryo-EM reconstruction have been deposited in the Protein Data Bank and EM data bank with the accession codes 9IVD and EMD-60925, respectively. The mass spectrometry proteomics data have been deposited to the ProteomeXchange Consortium via the PRIDE partner repository with the dataset identifier PXDxxxxxx.

---

**Fig. S1.**
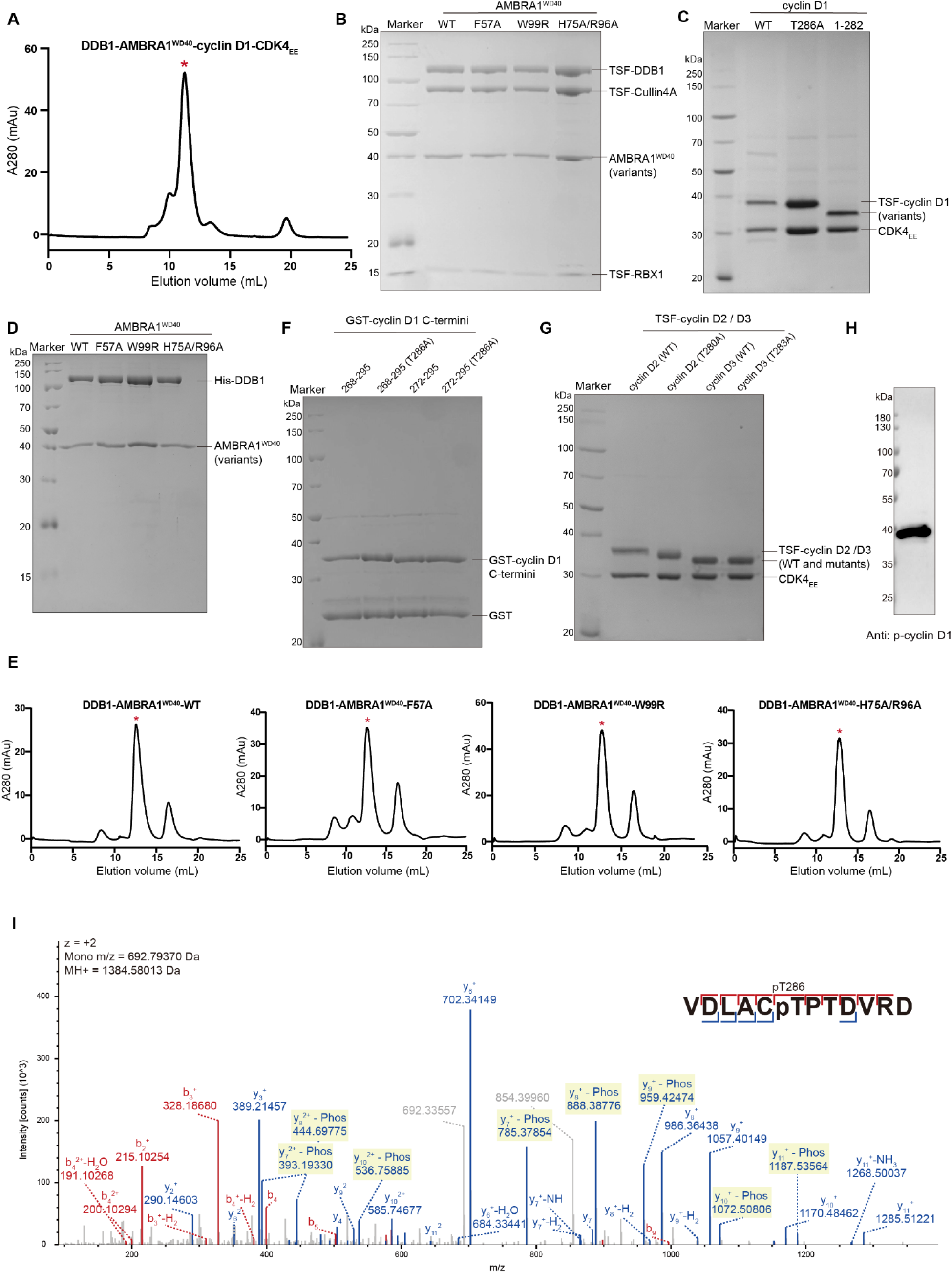
The SDS-PAGE and gel filtration profile of purified protein. (A) The gel filtration profile of the DDB1-AMBRA1^WD40^-cyclin D1-CDK4_EE_, the peak was used for cryo-EM structure analysis. (B) The SDS-PAGE of purified AMBRA1^WD40^-WT or mutants in complex with E3 ligase. (C) The SDS-PAGE of purified cyclin D1 WT, mutant and truncation in complex with CDK4_EE_. (D) The SDS-PAGE of purified DDB1-AMBRA1^WD40^-WT or mutants complexes, and corresponded gel filtration profiles (**E**). **(F)** The SDS-PAGE of purified GST-cyclin D1 C-termini. **(G)** The SDS-PAGE of purified TSF-cyclin D2, TSF-cyclin D3 and mutants in complex with CDK4_EE_. **(H)** The phosphorylation of cyclin D1 at Thr286 was analysed by western blotting in the reconstituted DDB1-AMBRA1^WD40^-cyclin D1-CDK4_EE_ protein complex. **(I)** MS/MS spectra for the identified phosphorylation site Thr286 in cyclin D1 from purified DDB1-AMBRA1^WD40^-cyclin D1-CDK4_EE_ protein complex.

**Fig. S2.**
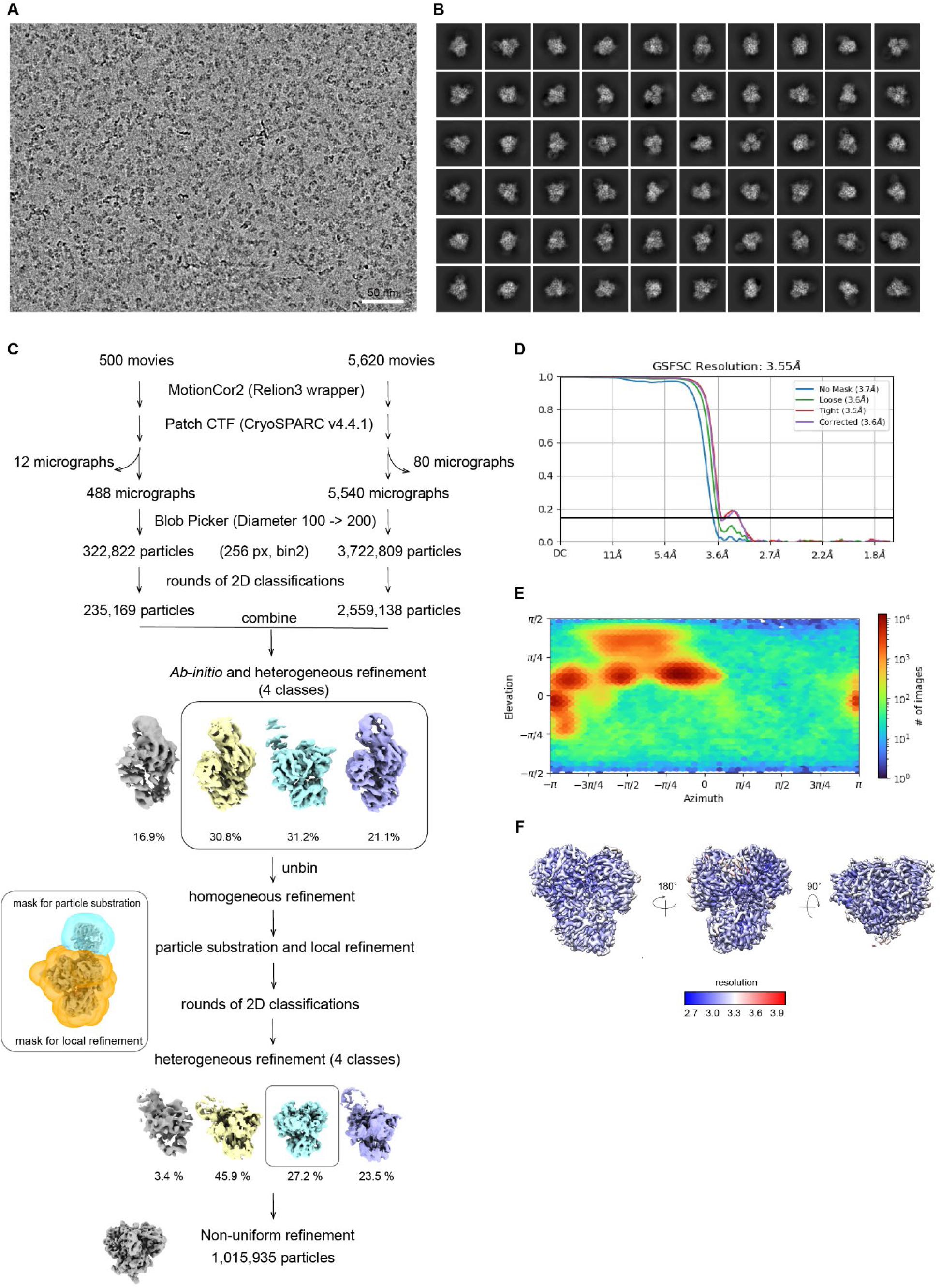
Cryo-EM data process of the DDB1-AMBRA1^WD40^-cyclin D1 complex. **(A)** A representative motion-corrected cryo-EM micrograph of the DDB1-AMBRA1 ^WD40^-cyclin D1 complex. Scale bar equals 50 nm. **(B)** Representative 2D class averages for the DDB1-AMBRA1^WD40^-cyclin D1 complex. **(C)** Flow chart of cryo-EM data processing. **(D)** The FSC plots are between two independently refined half-maps with no mask (blue), loose mask (green), tight mask (red), and corrected (purple). A cut-off of 0.143 (blue line) was used to estimate the resolution. **(E)** Angular particle distribution calculated in cryoSPARC for particle projections. The heatmap shows the number of particles for each viewing angle. **(F)** Local resolution is colored as indicated in the scale.

**Fig. S3.**
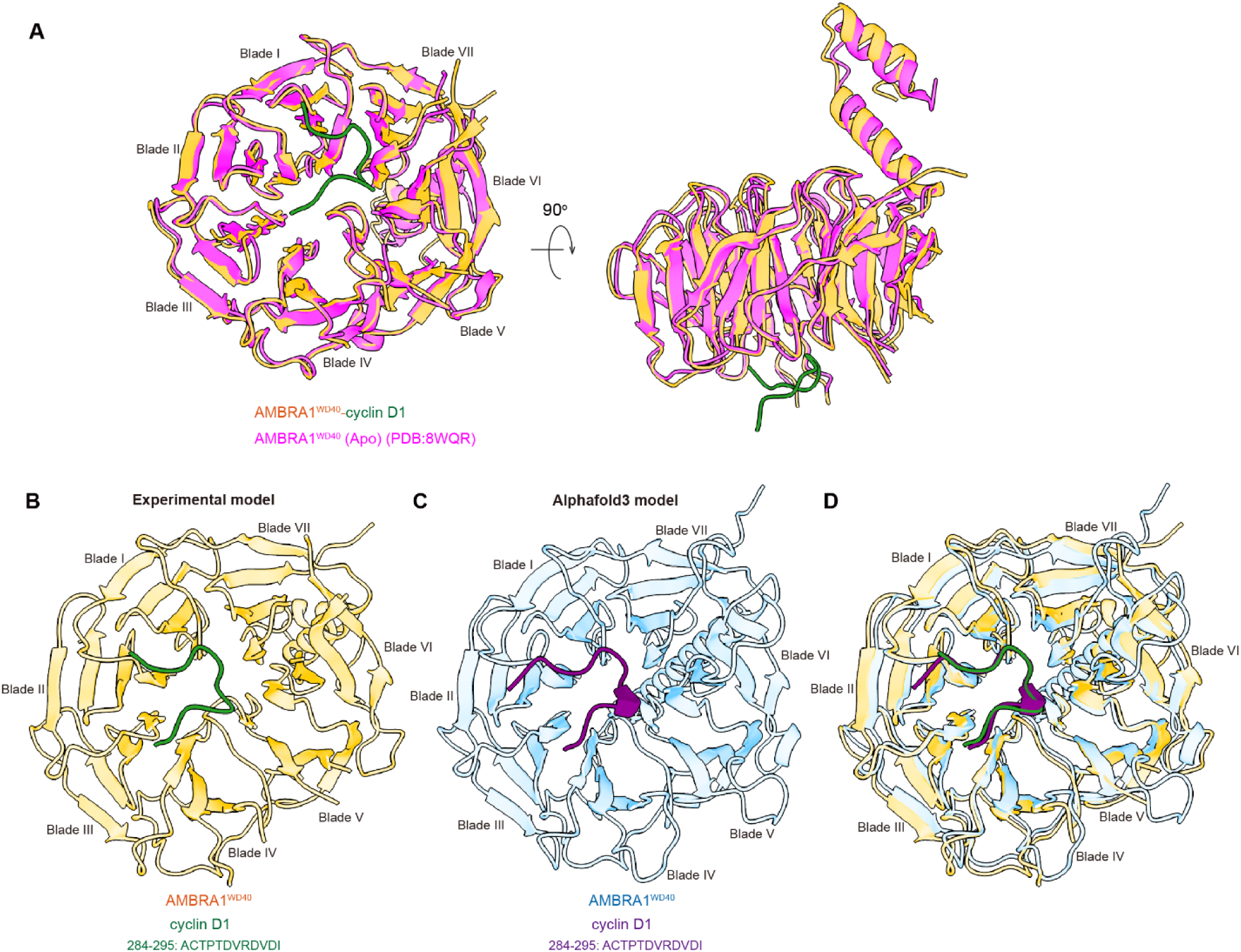
The comparison between experimental and AlphaFold3 model of AMBRA1^WD40^- cyclin D1. **(A)** The overlay of AMBRA1^WD40^-cyclin D1 and AMBRA1^WD40^ (Apo) (PDB:8WQR). The overall RMSD value between the AMBRA1^WD40^ models is 1.322 Å. AMBRA1^WD40^ colored in orange, cyclin D1 C-terminus colored in green and AMBRA1^WD40^ (Apo) colored in magenta. **(B)** The experimental model of AMBRA1^WD40^-cyclin D1 C-terminus (284-295). AMBRA1^WD40^ colored in orange and cyclin D1 C-terminus colored in green. **(C)** The predicted AlphaFold3 model of AMBRA1^WD40^-cyclin D1 C-terminus. AMBRA1^WD40^ colored in blue and cyclin D1 C-terminus colored in purple. **(D)** The overlay of experimental (**A**) and AlphaFold3 (**B**) model of AMBRA1^WD40^-cyclin D1. The RMSD value between the cyclin D1 C-terminus of experimental and predicted models is 0.324 Å.

**Fig. S4.**
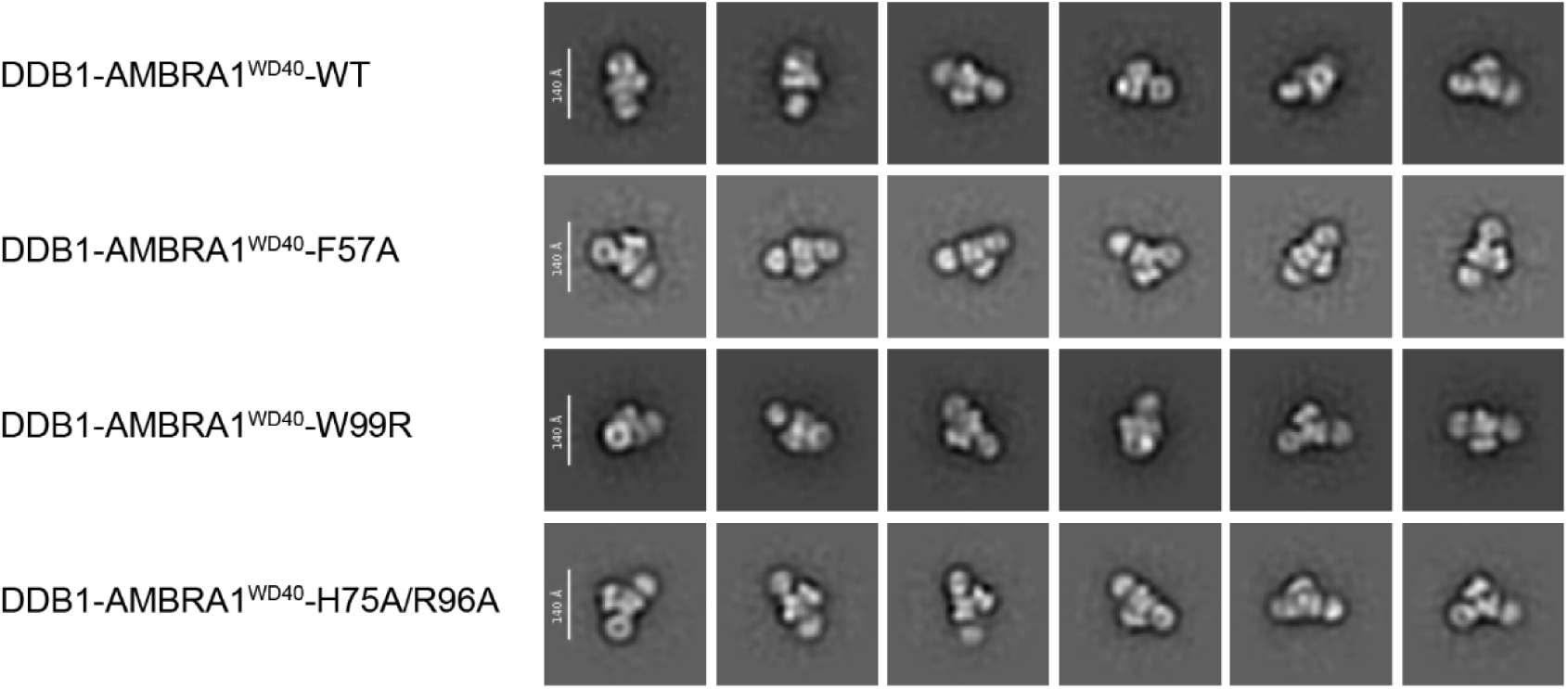
The representative 2D class averages of DDB1-AMBRA1^WD40^ complex. The representative 2D class average of negative staining shows the complete WD40 domain of AMBRA1^WD40^ mutants.

**Fig. S5.**
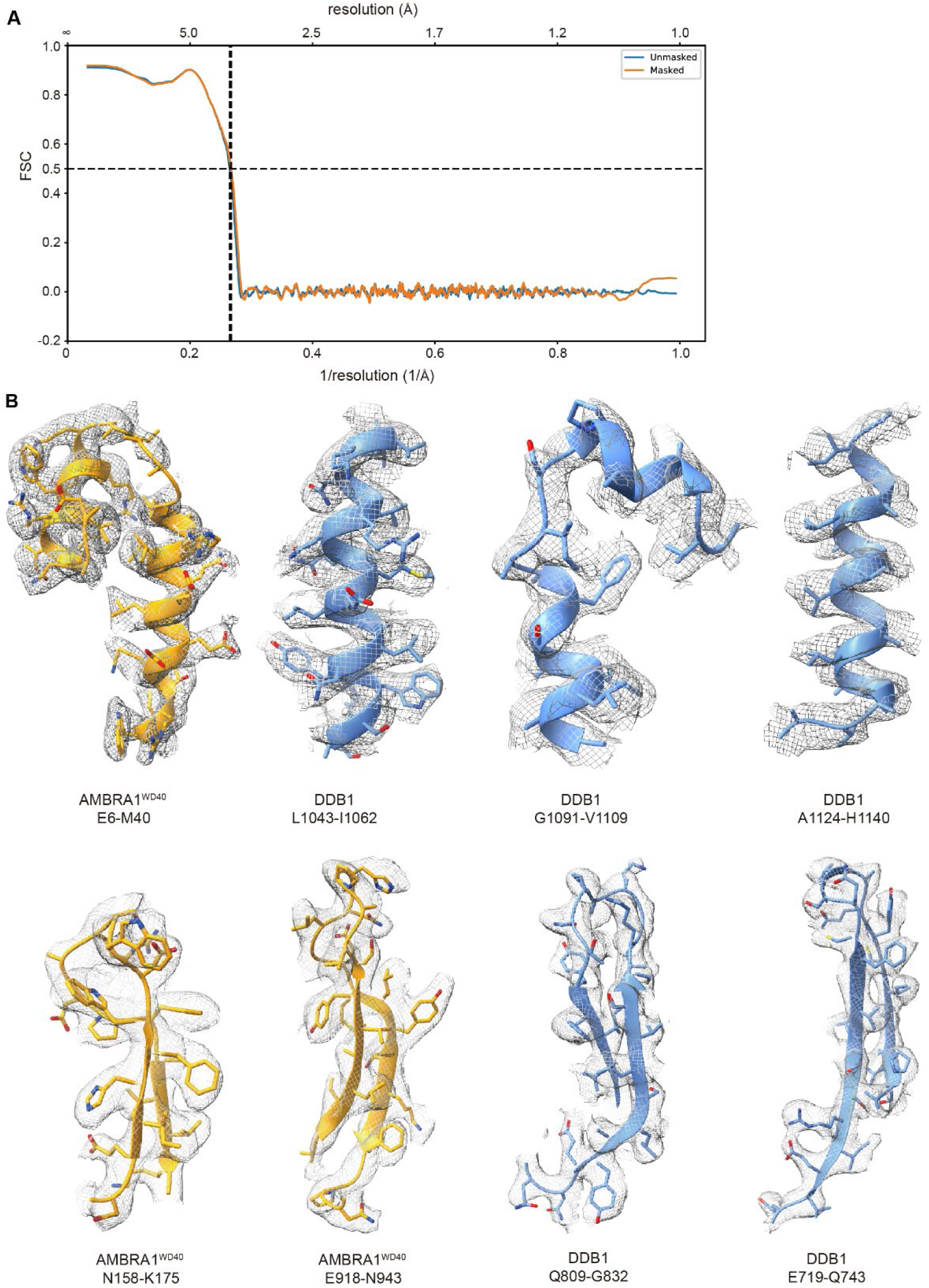
Model to map fitting. **(A)** FSC between the model and map for the DDB1-AMBRA1^WD40^-cyclin D1 complex against the cryo-EM map. **(B)** Representative cryo-EM densities fitted to the model.

**Fig. S6.**
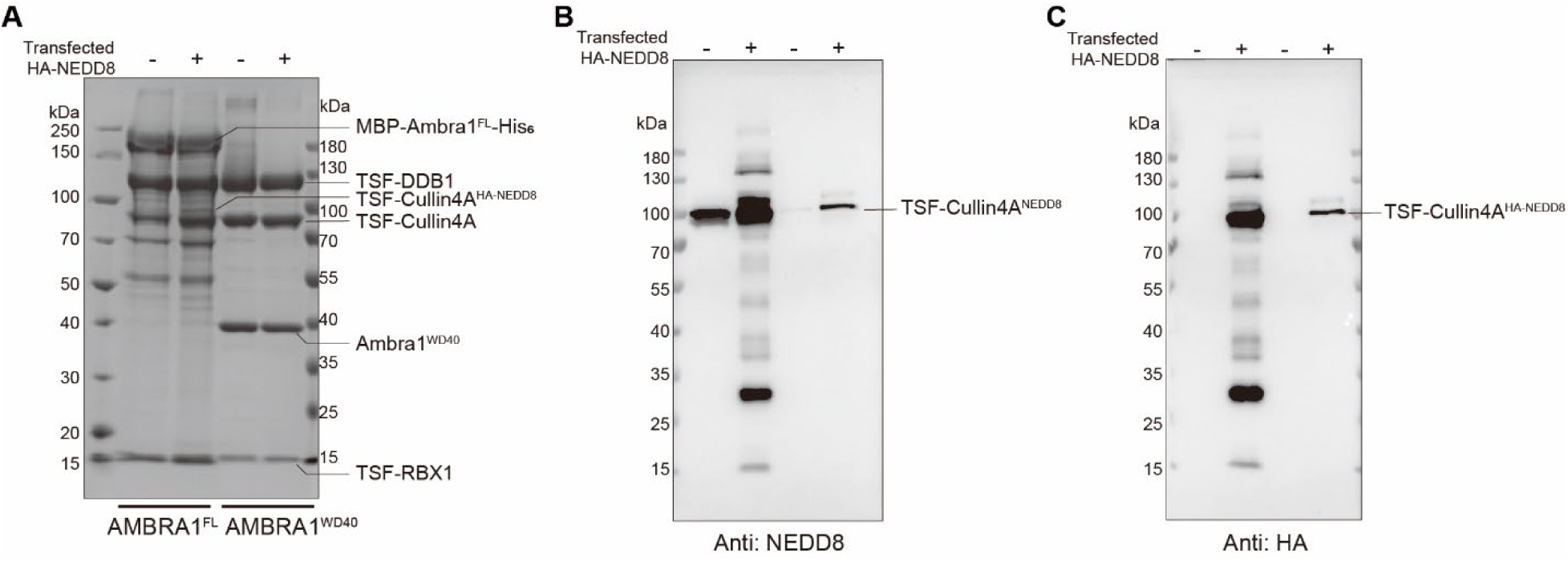
The SDS-PAGE and western blotting of the purified CRL4^AMBRA1 FL / WD40^ complex. **(A)** The SDS-PAGE of the purified Cullin4A-DDB1-RBX1-AMBRA1^WD40/FL^ complex. The complexes were expressed without and with HA-NEDD8. **(B)** The western blotting of purified Cullin4A-DDB1-RBX1-AMBRA1^FL/WD40^ complex corresponding to **(A)**, and detected by anti-NEDD8 antibody **(B)** and anti-HA antibody **(C)**.

**Table S1.**
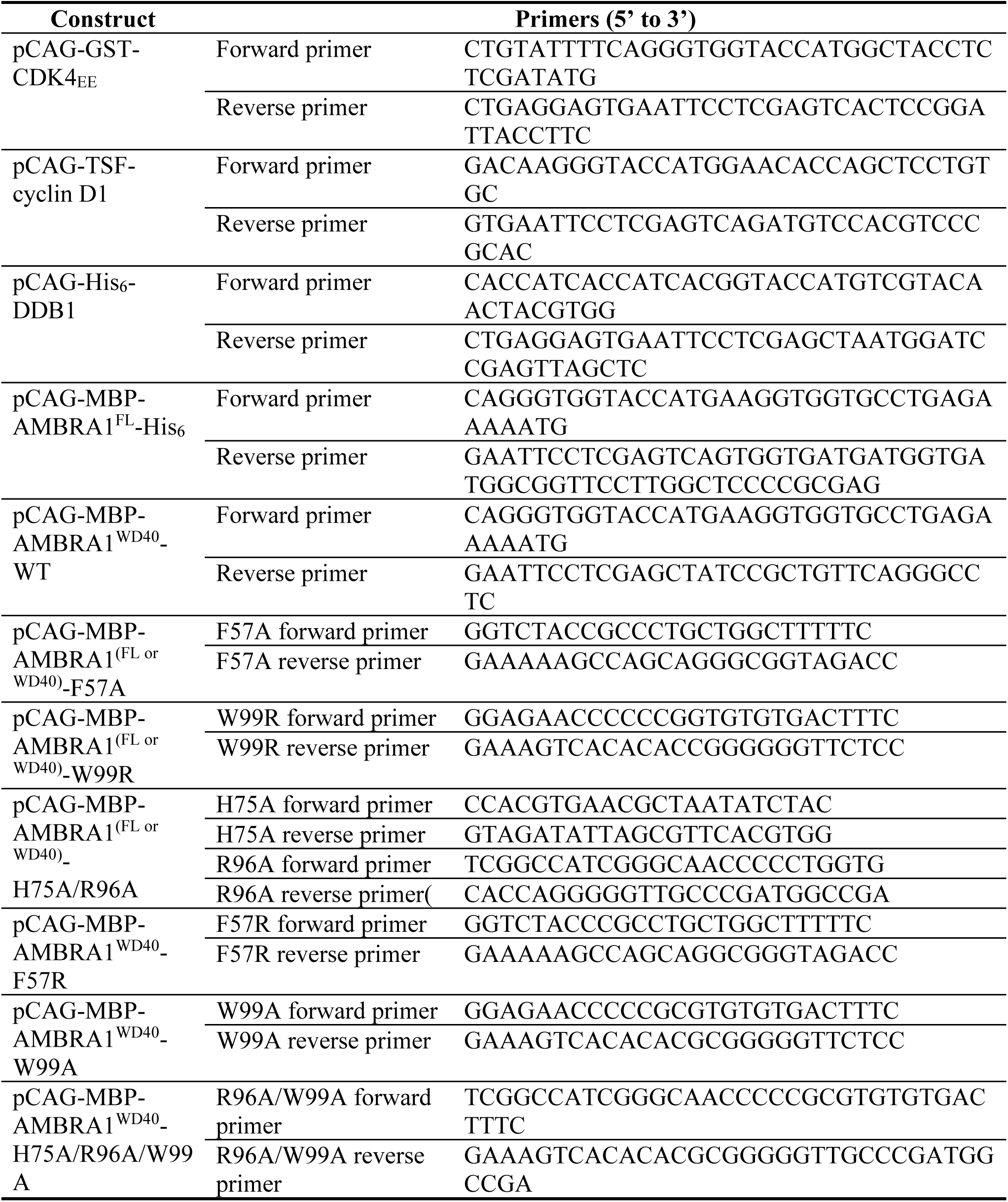
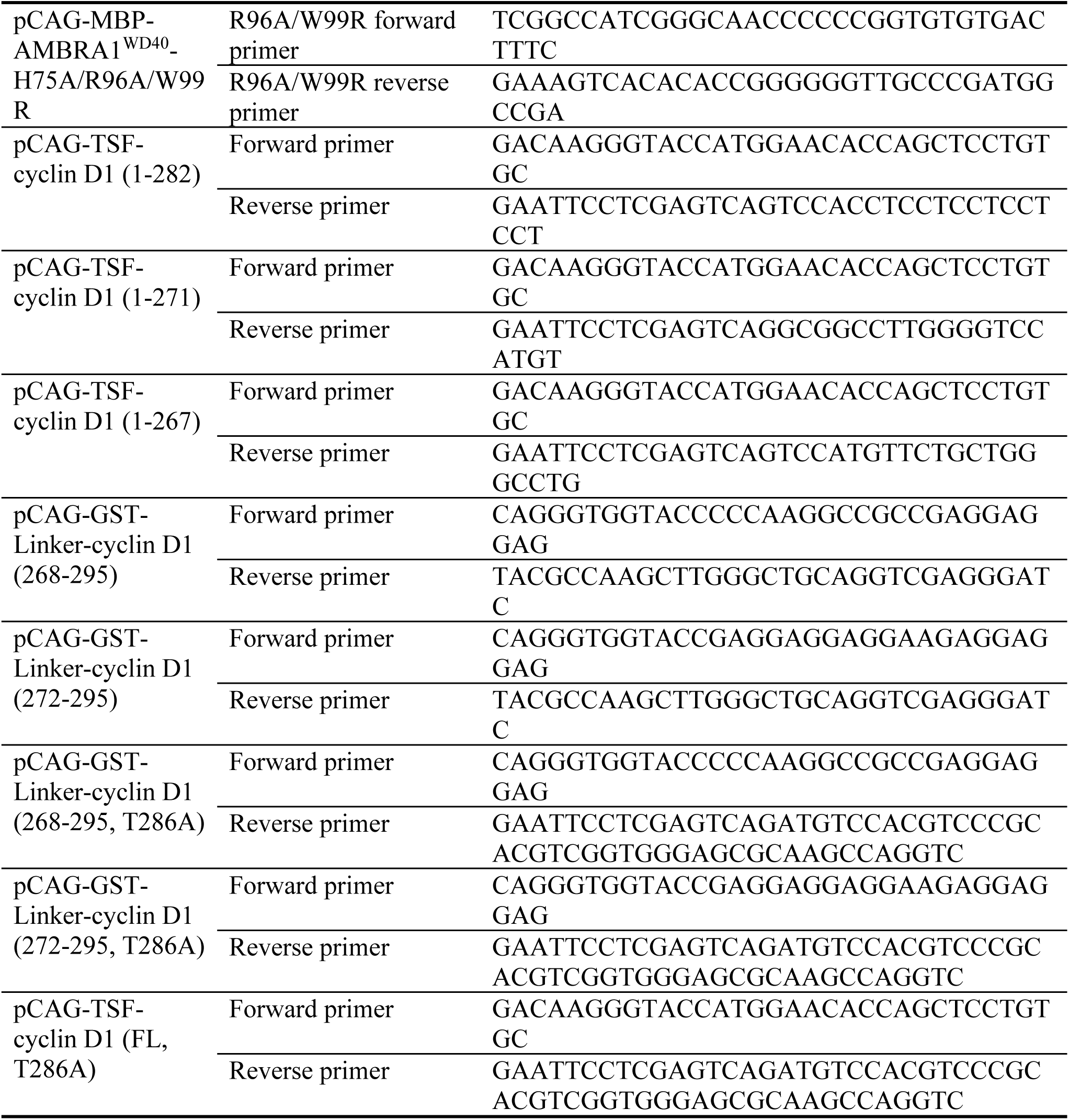
Primers used in this study.

**Table S2.**
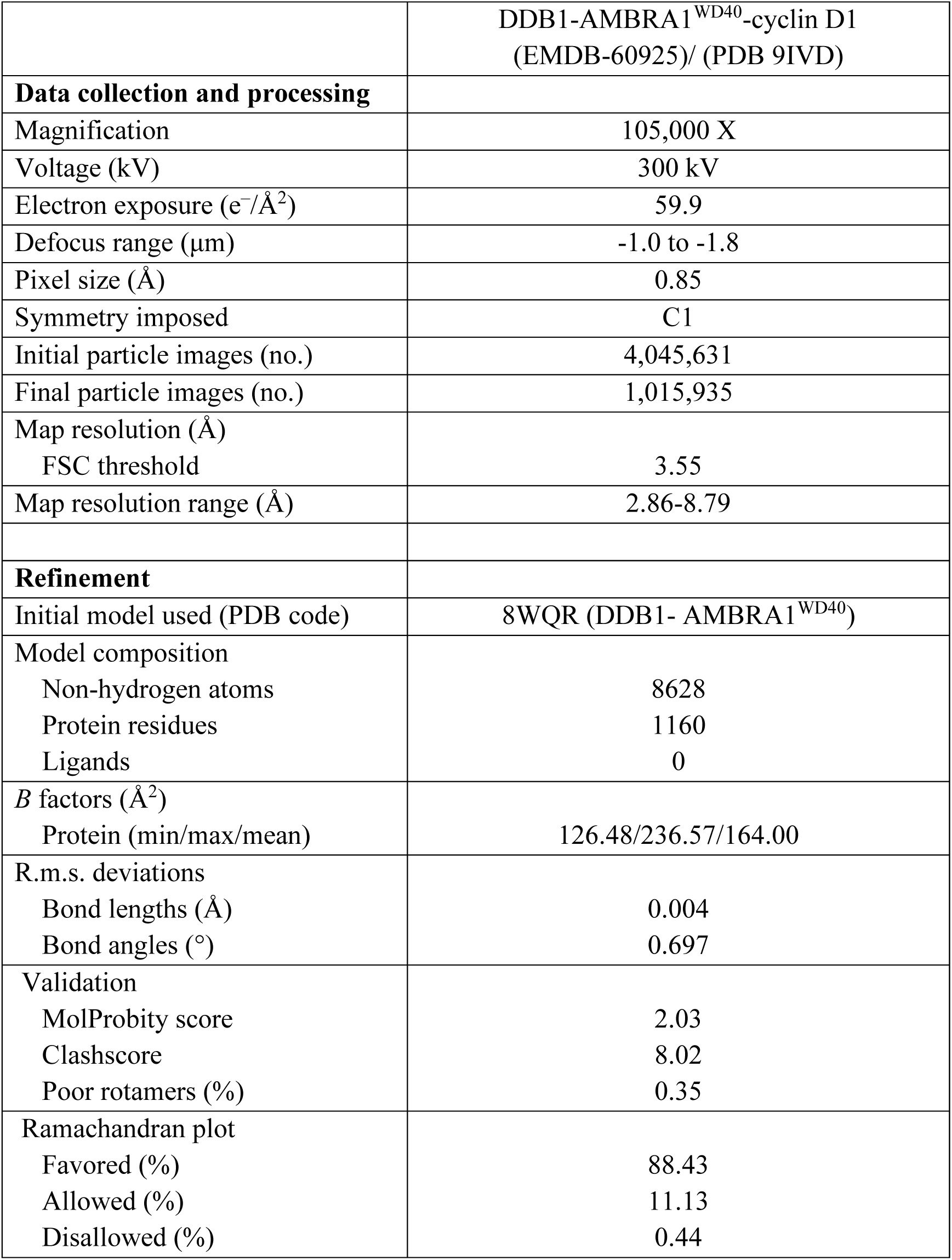
Cryo-EM data collection, refinement and validation statistics.

|  |  |
| --- | --- |
|  | DDB1-AMBRA1<sup>WD40</sup>-cyclin D1<br>(EMDB-60925)/ (PDB 9IVD) |
| **Data collection and processing** |  |
| Magnification | 105,000 X |
| Voltage (kV) | 300 kV |
| Electron exposure (e<sup>-</sup>/Å<sup>2</sup>) | 59.9 |
| Defocus range (μm) | -1.0 to -1.8 |
| Pixel size (Å) | 0.85 |
| Symmetry imposed | C1 |
| Initial particle images (no.) | 4,045,631 |
| Final particle images (no.) | 1,015,935 |
| Map resolution (Å) |  |
| FSC threshold | 3.55 |
| Map resolution range (Å) | 2.86-8.79 |
| **Refinement** |  |
| Initial model used (PDB code) | 8WQR (DDB1- AMBRA1<sup>WD40</sup>) |
| Model composition |  |
| Non-hydrogen atoms | 8628 |
| Protein residues | 1160 |
| Ligands | 0 |
| *B* factors (Å<sup>2</sup>) |  |
| Protein (min/max/mean) | 126.48/236.57/164.00 |
| R.m.s. deviations |  |
| Bond lengths (Å) | 0.004 |
| Bond angles (°) | 0.697 |
| Validation |  |
| MolProbity score | 2.03 |
| Clashscore | 8.02 |
| Poor rotamers (%) | 0.35 |
| Ramachandran plot |  |
| Favored (%) | 88.43 |
| Allowed (%) | 11.13 |
| Disallowed (%) | 0.44 |

